# The anti-cancer compound JTE-607 reveals hidden sequence specificity of the mRNA 3′ processing machinery

**DOI:** 10.1101/2023.04.11.536453

**Authors:** Liang Liu, Angela M Yu, Xiuye Wang, Lindsey V. Soles, Yiling Chen, Yoseop Yoon, Kristianna S.K. Sarkan, Marielle Cárdenas Valdez, Johannes Linder, Ivan Marazzi, Zhaoxia Yu, Feng Qiao, Wei Li, Georg Seelig, Yongsheng Shi

**Affiliations:** Department of Microbiology and Molecular Genetics, School of Medicine, University of California, Irvine, Irvine, CA 92697, USA; Center for Virus Research, University of California, Irvine, Irvine, CA 92697, USA; Department of Electrical and Computer Engineering, University of Washington, Seattle, Seattle, WA 98195, USA; Department of Biological Chemistry, School of Medicine, University of California, Irvine, Irvine, CA 92697, USA; Department of Genetics, Stanford University, Stanford, CA 94305, USA; Department of Statistics, University of California, Irvine, Irvine, CA 92697, USA; Paul G Allen School of Computer Science and Engineering, University of Washington, Seattle, Seattle, WA 98195, USA; Present address: Guangzhou Laboratory, Guangzhou, Guangdong, 510005, China; These authors contributed equally: Liang Liu, Angela M Yu

**Keywords:** mRNA 3′ processing, cleavage and polyadenylation, transcription termination, CPSF73, JTE-607, cancer, myeloid leukemia, machine learning

## Abstract

JTE-607 is a small molecule compound with anti-inflammation and anti-cancer activities. Upon entering the cell, it is hydrolyzed to Compound 2, which directly binds to and inhibits CPSF73, the endonuclease for the cleavage step in pre-mRNA 3′ processing. Although CPSF73 is universally required for mRNA 3′ end formation, we have unexpectedly found that Compound 2- mediated inhibition of pre-mRNA 3′ processing is sequence-specific and that the sequences flanking the cleavage site (CS) are a major determinant for drug sensitivity. By using massively parallel in vitro assays, we have measured the Compound 2 sensitivities of over 260,000 sequence variants and identified key sequence features that determine drug sensitivity. A machine learning model trained on these data can predict the impact of JTE-607 on poly(A) site (PAS) selection and transcription termination genome-wide. We propose a biochemical model in which CPSF73 and other mRNA 3′ processing factors bind to RNA of the CS region in a sequence-specific manner and the affinity of such interaction determines the Compound 2 sensitivity of a PAS. As the Compound 2-resistant CS sequences, characterized by U/A-rich motifs, are prevalent in PASs from yeast to human, the CS region sequence may have more fundamental functions beyond determining drug resistance. Together, our study not only characterized the mechanism of action of a compound with clinical implications, but also revealed a previously unknown and evolutionarily conserved sequence-specificity of the mRNA 3′ processing machinery.

## Introduction

Almost all eukaryotic mRNA 3′ end are formed through two chemical reactions, an endonucleolytic cleavage followed by polyadenylation^1,2^. Pre-mRNA 3′ processing is not only an essential step in gene expression, but also an important mechanism for gene regulation. ∼70% of human genes produce multiple mRNA isoforms by selecting different poly(A) sites (PASs), a phenomenon called alternative polyadenylation (APA)^3–5^. Distinct APA isoforms from the same gene can produce functionally different proteins and/or they can be regulated differently. APA is regulated in a developmental stage-and tissue-specific manner and mis-regulation of APA contributes to many human diseases. It remains poorly understood how APA is regulated in physiological or pathological contexts and pharmacological tools are needed for manipulating APA for research and therapeutic purposes.

The sites for canonical mRNA 3′ processing, or PASs, are defined by several cis-elements, including the AAUAAA hexamer, the U/GU-rich downstream elements, and other auxiliary sequences^1,2^. These cis-elements are recognized by multiple trans acting factors, including cleavage and polyadenylation specificity factor (CPSF) and cleavage stimulation factor (CstF), which in turn recruit other mRNA 3′ processing factors to assemble the pre-mRNA 3′ processing complex. Pre-mRNA cleavage is carried out by the endonuclease CPSF73^6^, which, together with CPSF100 and symplekin, forms the nuclease module of the CPSF complex mCF^7^. CPSF73 preferentially cleaves after CA or UA sequences^8^. Although the sequences flanking the CS display distinct and well-conserved nucleotide composition patterns^9–11^. it remains unknown what role, if any, these sequences play in pre-mRNA 3′ processing.

In recent years, CPSF73 has emerged as a drug target for treating a variety of diseases. For example, a number of small molecule drugs for treating toxoplasma gondii (causes toxoplasmosis)^12^, African trypanosomes (causes sleeping sickness)^13^, and Plasmodium (causes malaria)^14^, target the CPSF73 homologues in these pathogens. JTE-607 is a small molecule that inhibits the production of multiple cytokines by mammalian cells^15–17^. Animal studies demonstrated that administration of JTE-607 results in improvements in several inflammation diseases, including septic shock, acute injury, and endotoxemia^15–17^. Furthermore, JTE-607 was recently shown to have anti-cancer activities and specifically kill myeloid leukemia and Ewing’s sarcoma cells^18,19^. JTE-607 is a prodrug and is hydrolyzed to Compound 2 upon entering the cells by the cellular enzyme CES1^18,19^. Compound 2 specifically binds to CPSF73 near its active site to inhibit its activity^18^. In addition to its potential clinical application, JTE-607 has quickly become an important tool for research^20,21^. However, it is unclear if all pre-mRNA 3′ processing events in the human transcriptome are equally affected by JTE-607 and it is unclear why this compound is only active against specific cancer types.

Although the JTE-607 target, CPSF73, is universally required for pre-mRNA 3′ processing, we have found, surprisingly, that JTE-607-mediated inhibition of pre-mRNA 3′ processing is sequence-specific both in vitro and in cells. We have identified the CS region as a major determinant of drug sensitivity. Using massively parallel in vitro assay (MPIVA) coupled with machine learning, we have comprehensively characterized the relationship between the CS sequence and JTE-607 sensitivity and identified key sequence features that determine drug sensitivity. Using the MPIVA data, we trained a machine learning model, C3PO, that can accurately predict JTE-607 sensitivity of a PAS based on its CS region sequence. We demonstrated that C3PO can predict the effect of JTE-607 on PAS selection and transcription termination genome-wide. Together, our study not only better characterized the properties of an anti-cancer and anti-inflammation compound, but also revealed a previously unknown sequence-specificity of the mRNA 3′ processing machinery.

## Results

### Compound 2-mediated inhibition of mRNA 3′ processing in vitro is sequence-dependent

To better understand the mechanism of action for Compound 2, the active form of JTE-607^18^, we characterized its effect on pre-mRNA processing in an in vitro cleavage assay using HeLa cell nuclear extract (NE). We first performed in vitro cleavage assays with L3, the PAS of the adenovirus major late transcript, in the presence of DMSO or increasing concentration of Compound 2 (0.1, 0.5, 2.5, 12.5, 62.5, and 100 µM). Our results showed that the cleavage of L3 PAS was strongly inhibited by Compound 2 with an IC_50_ (concentration needed to achieve 50% of maximal inhibition) of 0.8 µM (Fig. 1a). Compound 2-mediated inhibition of pre-mRNA cleavage could occur at the cleavage step and/or the earlier pre-mRNA 3′ processing complex assembly step. To distinguish between these possibilities, we monitored pre-mRNA 3′ processing complex assembly on L3 PAS in the presence of DMSO or increasing concentrations of Compound 2 using an electrophoretic mobility shift assay. The pre-mRNA 3′ processing complex assembled indistinguishably under all conditions tested (Fig. 1b). These results suggest that Compound 2 does not interfere with pre-mRNA 3′ processing complex assembly, but blocks cleavage of the L3 PAS.

**Fig 1.**
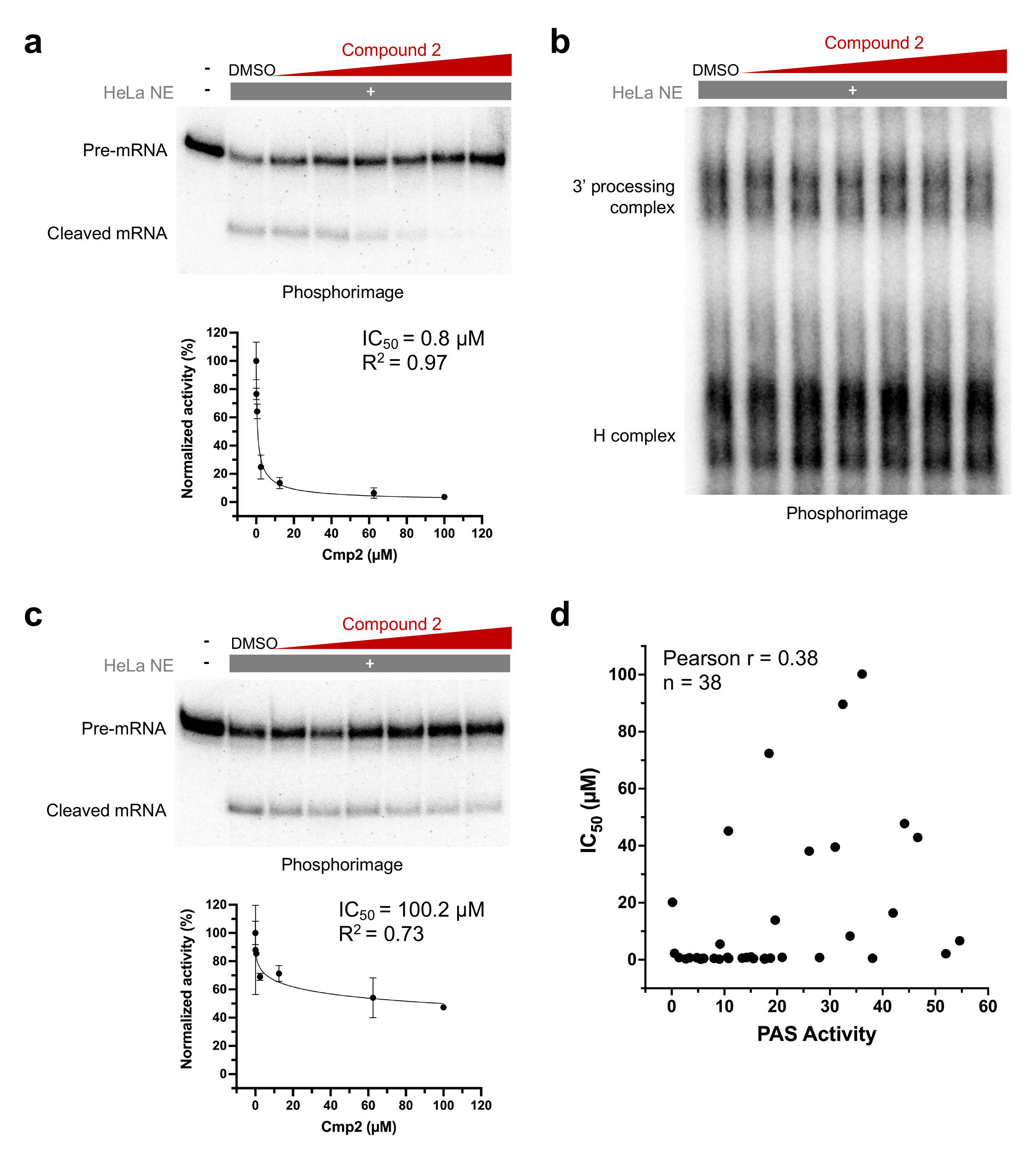
Compound 2-mediated inhibition of mRNA 3’ processing in vitro is sequence-dependent. **(a)** In vitro cleavage assay on L3 PAS with increasing concentration of Compound 2 and its IC_50_ quantification. Radio-labeled RNAs from the reactions were extracted and resolved on 8M urea gel and visualized by phosphorimaging. Compound 2 concentrations used are: 0.1, 0.5, 2.5, 12.5, 62.5, and 100 µM. **(b)** Electrophoretic mobility shift assay (EMSA) with L3 PAS in the presence of increasing concentration of Compound 2. Same concentrations as (A) were used. **(c)** In vitro cleavage assay on SVL PAS with increasing concentration of Compound 2 and its IC_50_ quantification. **(d)** PAS activity and IC_50_ correlation of 34 in vitro tested PAS.

We next performed similar in vitro cleavage assays on other PASs. Surprisingly, we found that different PASs displayed different sensitivities to Compound 2-mediated inhibition of pre-mRNA cleavage. For example, significant cleavage was observed for SVL, the PAS from SV40 late transcript, even at the highest concentration tested of Compound 2 (Fig. 1c). The estimated IC_50_ for SVL PAS was greater than 100.2 μM (Fig. 1c). Therefore, the IC_50_ of L3 and SVL PASs differ by over 100-fold. Similar to L3, mRNA 3′ processing complex assembly on SVL PAS was not affected by Compound 2 (Fig. S1). In total we performed the same in vitro cleavage assays with 38 different PASs and found that their IC_50_ values varied widely (Fig. 1d). To begin to understand the molecular basis for such variations, we first asked if the Compound 2 sensitivity of a PAS is determined by its strength, i.e. the efficiency by which it is processed by the pre-mRNA 3′ processing machinery. We measured the percentage of pre-mRNA cleaved in vitro in the absence of the drug and compared this value with their IC_50_. Our results detected poor correlation between the two measurements (r=0.38) (Fig. 1d, SI Table 1). We conclude that the cleavage of different PASs display differential sensitivity to Compound 2 in vitro and that the sensitivity of a PAS is not determined by its strength.

### The cleavage site (CS) region sequence is a major determinant of Compound 2 sensitivity

Since PASs display sequence-dependent sensitivity to Compound 2 in vitro, we next wanted to map the specific region(s) of the PAS that determine its drug sensitivity. To this end, we divided the PAS sequence into three regions: the AAUAAA hexamer and upstream sequence (referred to as upstream sequence or UPS), the CS region (20 nucleotide (nt) region centered at the cleavage site), and the downstream sequence (DS) (Fig. 2a). Among PAS sequences we tested previously, L3 (IC_50_=0.8 μM, Fig. 1a) and SVL (IC_50_=100.2 μM, Fig. 1c) showed the lowest and the highest resistance to Compound 2, respectively. Therefore, we constructed a series of chimeric PASs between these sequences, in which one or more of the three regions in one PAS was replaced by their counterparts in another. We then measured their IC_50_ using in vitro cleavage assay as described above. Replacing the UPS of L3 PAS with that of SVL did not result in a major change in IC_50_ (Chimera 1, IC_50_=2.1 μM, Fig. 2a and Fig. S2a). However, replacing both the UPS and the CS of L3 with those of SVL dramatically increased the resistance to Compound 2 (Chimera 2, IC_50_=89.6 μM, Fig. 2a and Fig. S2b), suggesting that the CS region plays a major role. On the other hand, replacing the UPS of SVL with that of L3 led to a significant decrease in drug resistance (Chimera 3, Fig. 2a and Fig. S2c), although its IC_50_ (39.5 μM) was still nearly 50 times higher than that of L3. Replacing both the UPS and CS of SVL with those of L3 led to a near 15- fold decrease in IC_50_ (Chimera 4, IC_50_=6.7 μM) (Fig. 2a and Fig. S2d), again highlighting a major role for the CS region. By contrast, the DS did not seem to play a significant role (compare L3 and Chimera 4 or SVL and Chimera 2, Fig. 2a). Given the large impact of the CS region on Compound 2 sensitivity in both backgrounds, we swapped the CS regions alone between L3 and SVL. The results showed that replacing the L3 CS region with that of SVL increased its IC_50_ to 47.8 μM, a near 60-fold increase (L3-SVL CS, Fig. 2a and b). Even more dramatically, the opposite change in SVL reduced its IC_50_ to 0.8 μM (SVL-L3 CS, Fig. 2a and c), identical to that of L3. These results demonstrated that the CS region is a major determinant of Compound 2 sensitivity in both backgrounds. Additionally, the UPS also contributes to the drug sensitivity in a context-dependent manner, while the DS does not appear to play a significant role. Therefore, we have focused on the CS region for the rest of this study.

**Fig 2.**
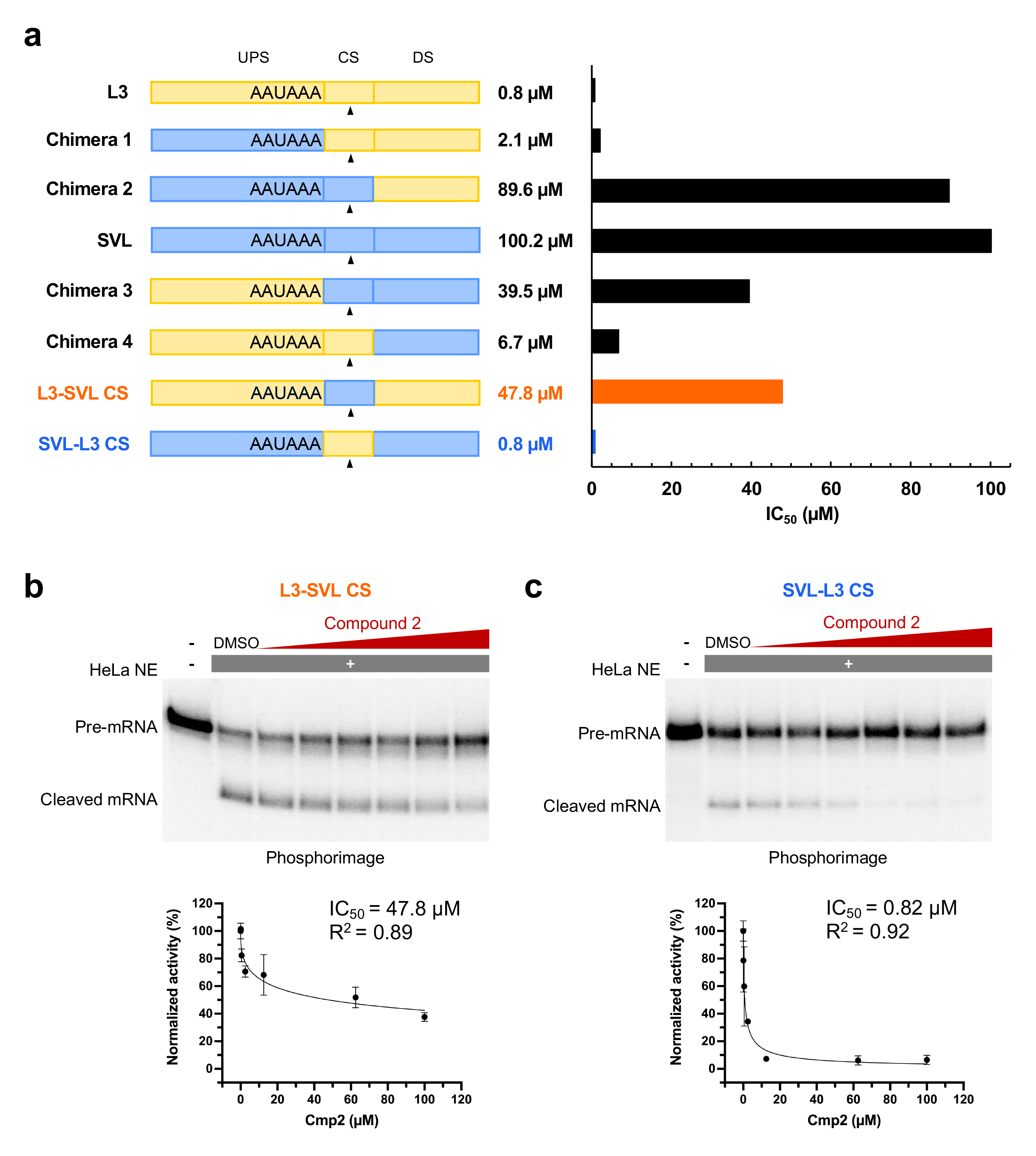
Cleavage site (CS) region is a major determinant of Compound 2 sensitivity. **(a)** A diagram of L3, SVL and their chimeras. Their corresponding IC_50_ were plotted on the right. UPS: upstream sequence; CS: cleavage site; DS: downstream sequence. Black triangles denote the cleavage position YA (Y is U or C). **(b)** In vitro cleavage of L3-SVL CS with increasing concentration of Compound 2, similar to Fig. 1A and C. **(c)** In vitro cleavage of SVL-L3 CS with increasing concentration of Compound 2.

### Define the CS sequence-Compound 2 sensitivity relationship by using massively parallel in vitro assay (MPIVA)

We next wanted to comprehensively define the relationship between the CS region sequence and Compound 2 sensitivity. To this end, we designed an MPIVA strategy (Fig. 3a). Using L3 (sensitive) or SVL (resistant) PAS as backbones, we replaced the original cleavage site with a YA sequence (Y is U or C), which is the preferred cleavage site for CPSF73, and randomized the 23nt flanking sequence. These two libraries, called L3-N23 and SVL-N23, contained ∼3 million PAS variants each and were transcribed into RNA. The PAS RNA pools were used for in vitro cleavage and polyadenylation assays in the presence of DMSO (control) or increasing concentrations of Compound 2, including low (0.5 μM), medium (2.5 μM), and high (12.5 μM) concentrations. As shown in Fig. 3b, the PAS RNA pool was efficiently cleaved in vitro in the presence of DMSO and the cleavage efficiency gradually decreased in the presence of increasing concentrations of Compound 2. The starting PAS RNA pool and the cleaved RNA pools under different conditions were subjected to high throughput sequencing using the Illumina platform (Fig. 2b). For each variant, a resistance score was calculated as the log ratio between its frequency in Compound 2- treated samples and that in DMSO-treated samples. As shown in Fig. 3c and Fig. S3a, the resistance scores of all variants were concentrated in a narrow peak centered at ∼0 at low Compound 2 concentration (L3: −0.04 ± 0.31; SVL: −0.05 ± 0.28) but diverged more at high inhibitor concentration (L3: −0.12 ± 0.53; SVL: −0.09 ± 0.43), suggesting that, as expected, drug sensitivities are better distinguished at higher drug concentrations. Furthermore, we compared the resistance scores of all variants and their cleavage efficiency (log ratio between the frequency of a PAS variant in Library 2 and that in Library 1) and found that there was no significant correlation (Fig. 3d and Fig. S3b), which was consistent with Fig. 1d. Thus, both our low throughput in vitro assays and high throughput screen results demonstrated that the Compound 2 sensitivity of a PAS is not dependent on its strength.

**Fig 3.**
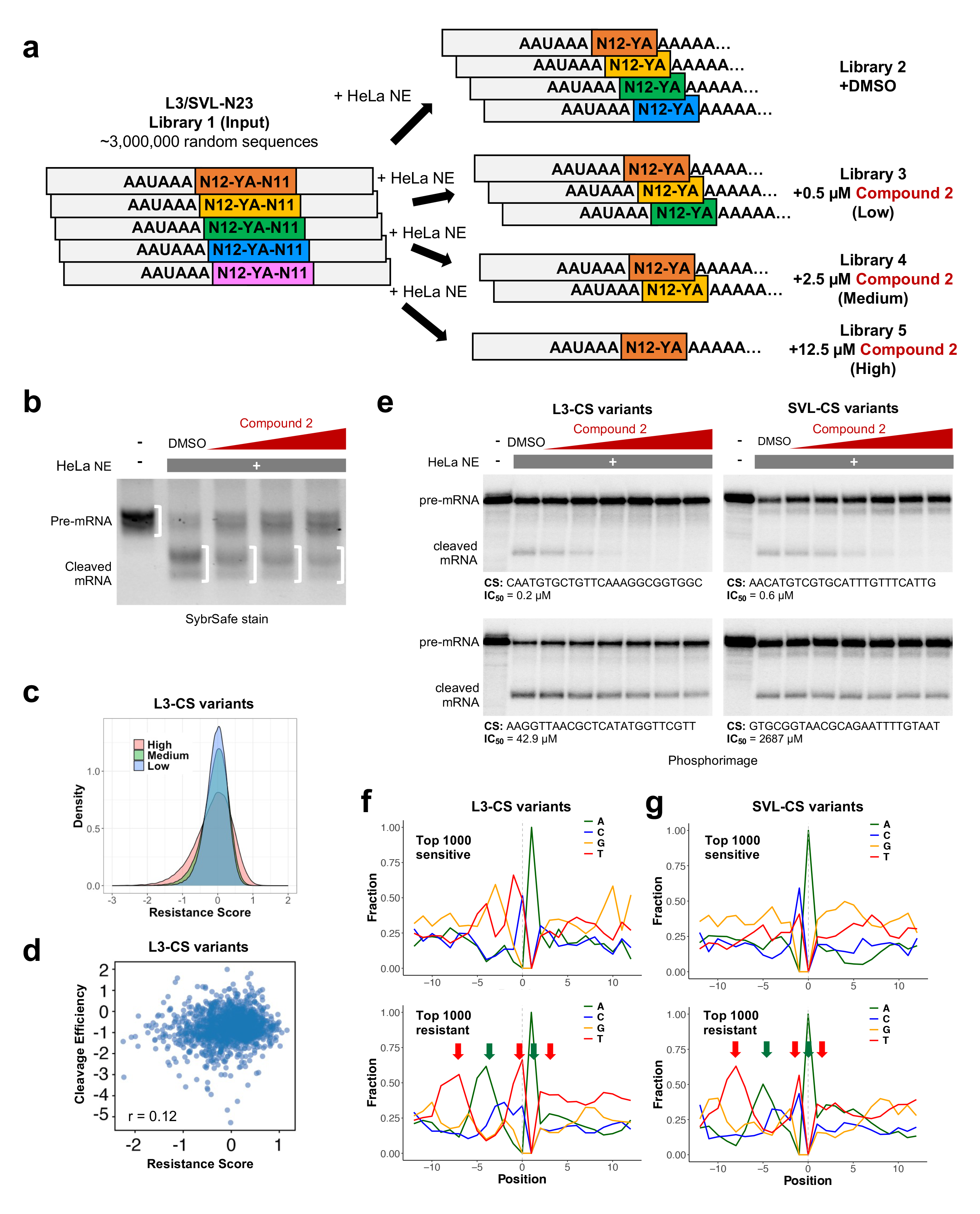
Determine sequence specificity for Compound 2 sensitivity by massively parallel in vitro assay (MPIVA). **(a)** Design of the randomized CS sequence libraries and the MPIVA assay. Each box represents a sequence variant. YA: cleavage position (Y is U or C). N: random nucleotide. **(b)** The randomized sequence library L3/SVL-N23 were transcribed into RNAs and used for in vitro cleavage/polyadenylation assays in the presence of 0.5, 2.5, and 12.5 µM Compound 2. The RNAs from these reactions were amplified by RT-PCR and resolved on an agarose gel. The RNA species were marked on the left. The white half brackets mark the regions on the gel that were extracted and amplified for sequencing. **(c)** A density plot for the resistance scores of all variants in L3-N23 library. The low, medium, and high groups represent the screens in the presence of 0.5, 2.5, and 12.5 μM Compound 2 as shown in (B). **(d)** A scatter plot comparing the cleavage efficiency log(frequency in Library 2/frequency in Library 1) and the resistance score (log(frequency in Library 5/frequency in Library 2) of L3-CS variants. Pearson correlation is shown. **(e)** Examples of validation experiments using in vitro cleavage assays for variants from both L3- and SVL-N23 libraries. **(f)** Nucleoside distribution of L3-CS variants for the top 1000 most sensitive and resistant sequences. **(g)** Nucleoside distribution of SVL-CS variants for the top 1000 most sensitive and resistant sequences. T-and A-rich regions were marked with red and green arrows respectively.

Based on the resistance scores in the high Compound 2 concentration condition, we obtained a list of the top 1000 most sensitive and resistant PASs from both the L3-N23 and SVL-N23 libraries. We selected 6 variants, 3 sensitive and 3 resistant, in each background and tested them using our in vitro cleavage assay and our data validated the screen results (Fig, 3e, Fig. S4). It was noted that some of the variants (e.g. Fig. 3e, top left panel) were more sensitive to Compound 2 than the original L3 while other variants displayed greater resistance than SVL (e.g. Fig. 3e, bottom right panel), indicating that our screens selected variants with a wide range of drug sensitivities. Interestingly, the nucleotide composition in the CS region of sensitive and resistant PASs showed distinct patterns. The CS regions of sensitive L3 variants are generally G/U-rich, especially in the region upstream of the cleavage site (Fig. 3f, top panel). By contrast, resistant CSs contained alternating U-rich and A-rich sequences mainly in the region upstream of the cleavage site (Fig. 3f, bottom panel). Very similar patterns were observed in SVL background (Fig. 3g), suggesting that the CS region sequence can determine Compound 2 sensitivity independent of other regions. Consistent with the nucleotide compositions, our motif analyses of the sensitive and resistant variants detected U/G-rich and A/U-rich motifs, respectively, in both L3 and SVL libraries (Fig. S5). These results defined the key sequence features in the CS region that determine Compound 2 sensitivity.

### Machine learning predictions of Compound 2 sensitivity from PAS sequences

We next used our MPIVA data to train a machine learning model with the goal of predicting Compound 2 sensitivity of any given PAS based on its CS region sequence. Our model, called Cleavage and Counteraction with Compound 2 on Polyadenylation Outcomes (C3PO), is a three-layer convolutional neural network (CNN) that is based on the Optimus 5′ architecture that we have previously used to predict polysome profiles from 5′ untranslated region (UTR) sequences (Fig. 4a, Methods).^22^ C3PO uses the 25 nt CS sequences as inputs and predicts Compound 2 sensitivity, which is calculated as the log ratio between each variant’s percent representation in the DMSO-treated and Compound 2-treated libraries (Fig. 3a). C3PO was trained on the processed MPIVA datasets from both the L3 and SVL RNA contexts, and model performance was assessed on held-out variants from both RNA contexts. We used the variants with high read coverage in the input and DMSO-treated data (Libraries 1 and 2) as our test set to minimize the impact of measurement noise (Methods). C3PO performed better on higher doses of Compound 2 with Pearson’s r of 0.56, 0.74, and 0.84 for 0.5 μM, 2.5 μM, and 12.5 μM, respectively. We explored variations of convolution-based machine learning architectures (SI Table 2), and this trend was consistent. This was expected as drug resistance is better detected at higher drug dose (Fig. 3c). Due to the better model performance at the highest dose of 12.5 μM, we focused further analyses on this regime.

**Fig 4.**
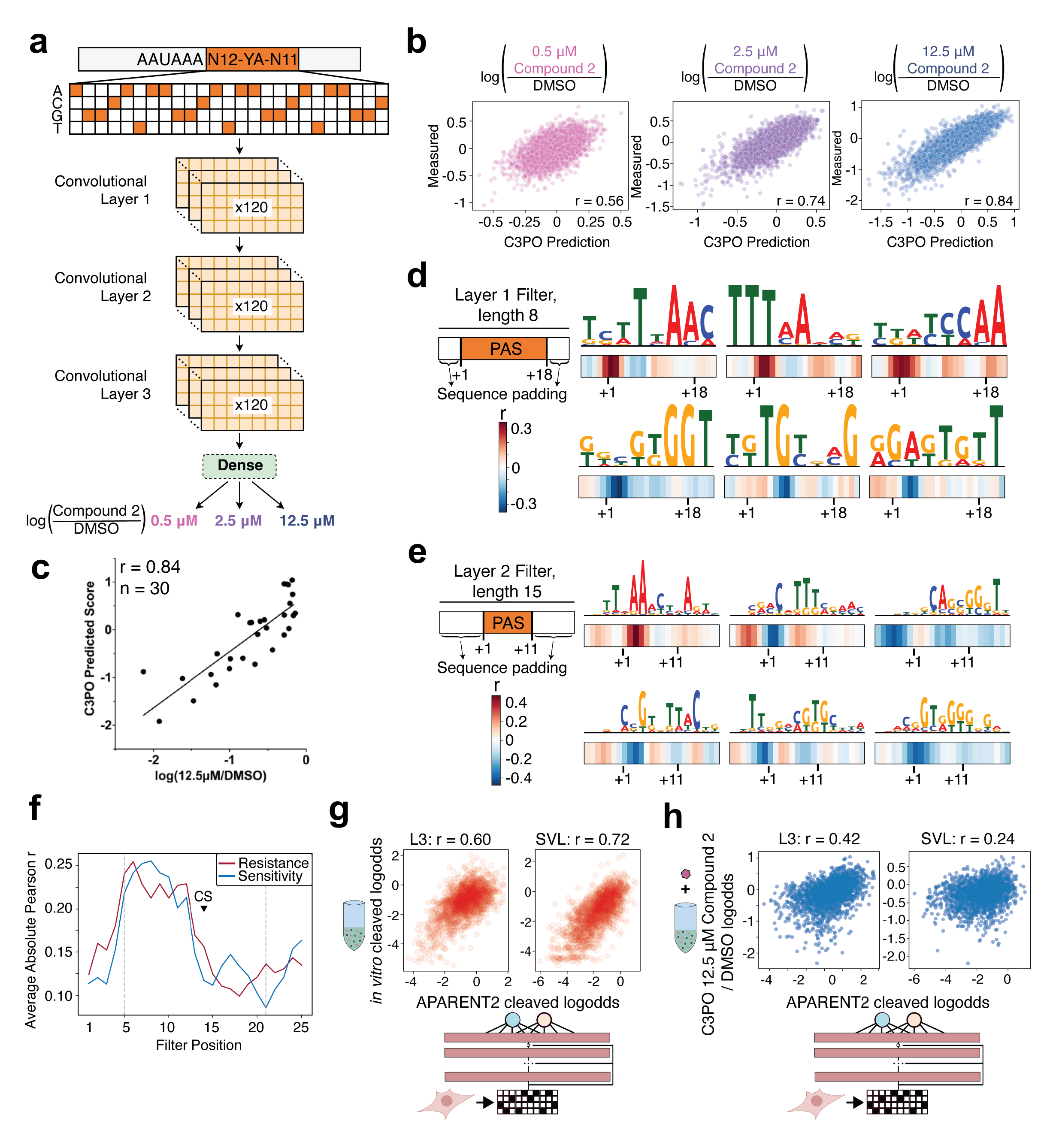
C3PO architecture, performance, and layer feature analyses. **(a)** The model takes 25 nt RNA sequences immediately downstream of the core hexamer and predicts three doses of Compound 2 drug sensitivity by predicting the log ratio of percent reads in a drug-treated sample to a DMSO-treated sample. **(b)** Scatter plots of C3PO performance on predicting drug sensitivity at 3 Compound 2 doses on test sequences. Test sequences include equal number of sequences derived from both the L3 and SVL RNA contexts. **(c)** A scatter plot comparing the resistance scores predicted by C3PO and those measured experimentally. **(d)** Convolutional layer 1 and **(e)** layer 2 max filter activations with the highest Pearson correlation with 12.5 μM Compound 2 predictions. Sequence logos are plotted on top of per-position absolute value of Pearson correlations with 12.5 μM Compound 2 sensitivity predictions. All layer 1 and 2 filters are reported in Fig. S6, S7. **(f)** Plot of average of all layer 1 filters’ absolute value of Pearson correlation with 12.5 μM Compound 2 predictions across all positions. These are split into Pearson correlation values associated with resistance or sensitivity. Dashed gray lines indicate positions at the edge of sequence padding. The position of the cleavage site (CS) is marked and note that preceding filters may overlap with the designed canonical cut sites. **(g)** Scatterplots of RNA cleavage logodds measured *in vitro* calculated from input and DMSO libraries versus those from APARENT2 predictions. **(h)** Scatterplots of Compound 2 resistance predicted by C3PO and the cleavage efficiency predicted by APARENT2.

To test the performance of C3PO, we compared the Compound 2 resistance scores (log(12.5 µM/DMSO)) of 30 distinct PASs measured by in vitro cleavage assays as shown in Fig. 1D (PASs that contain the same CS region sequences were omitted to avoid redundancy) and those predicted by C3PO. The C3PO predictions showed strong and positive correlation with experimental measurements with a Pearson r of 0.84 (Fig. 4c, SI Table 3). This is very similar to its performance on the MPIVA dataset (compare Fig. 4c with 4b, 12.5 µM panel). These results strongly suggest that C3PO can accurately predict Compound 2 sensitivity of PAS sequence in vitro.

We next wanted to identify sequence motifs that are predictive of Compound 2 sensitivity by extracting filter position weight matrices (Fig. 4a). The position-specific effect on Compound 2 sensitivity of each filter was quantified by measuring the correlation with drug sensitivity at each position across the CS region. Filters associated with higher resistance (dark red color) learned motifs that were A/U-rich, while lower resistance filters (dark blue) typically learned motifs with higher G/U content (Fig. 4d). Sequence motifs strongly associated with Compound 2 sensitivity predictions are positioned such that they begin upstream of the CS (Fig. 4d-f, Fig. S6). Layer 2 filters learn to use combinations of Layer 1 filters for predictions of drug sensitivity. 15-mers learned by layer 2 filters also showed A/U-rich and G/U-rich motifs for resistant and sensitive PASs respectively (Fig. 4e and Fig. S7). Interestingly, both resistance-and sensitivity-associated motifs are enriched in the region upstream of the CS (Fig. 4f).

Given the known function of RNA secondary structures in pre-mRNA 3′ processing^23^, we investigated the its potential impact on Compound 2 sensitivity. We compared the minimum free energy (MFE) structures for the top 10,000 resistant and sensitive sequences (Fig. S8a-b). The differences between ι1Gs for the resistant and sensitive sequences were modest, but statistically significant with p-values of < 2.2 x 10^-308^ and 1.58 x 10^-26^ and for L3 and SVL, respectively. The difference between base pairing probabilities for resistant and sensitive sequences also show different global patterns between the L3 and SVL backbones, indicating that background-specific secondary structural features may contribute to drug sensitivity (Fig. S8c-d). Taken together with C3PO’s ability to accurately predict Compound 2 sensitivity with sequence alone, our results suggest that sequence is the primary determinant of Compound 2 sensitivity while secondary structure may play a minor role.

We further explored the usage of machine learning models to characterize Compound 2 sensitivity and its relationship with processing efficiency. First, we compared the cleavage efficiency measured by our MPIVA assays with that predicted by APARENT2 ^24^, a highly accurate deep learning model for predicting cleavage/polyadenylation efficiency that was trained using massively parallel reporter assays in mammalian cells. We saw good correlation between APARENT2-predicted cleavage efficiency and our MPIVA data with a Pearson r of 0.60 for the L3 background and 0.72 for the SVL background (Fig. 4g). These results suggest that the CS region sequence can have a significant impact on cleavage efficiency, and that the cleavage efficiency values measured by our MPIVA system are highly consistent with measurements obtained in cells. Finally, we compared the resistance score predicted by C3PO with the cleavage efficiency predicted by APARENT2 for all CS variants and observed poor correlation with Pearson r = 0.42 and 0.24 for L3 and SVL respectively (Fig. 4h). This is consistent with our in vitro cleavage assay (Fig. 1d) and MPIVA results (Fig. 3d and Fig. S3b) and provided further evidence that the Compound 2 sensitivity of a PAS is not dependent on its strength.

### JTE-607 modulates PAS selection and transcription termination in a sequence-specific manner in human cells

To determine whether the sequence-specific sensitivity to Compound 2 observed in vitro was true in cells, we performed two genome-wide analyses. First, we analyzed the global APA profiles in DMSO-and JTE-607-treated human HepG2 cells using PAS-seq, a high throughput RNA 3′ sequencing method for quantitatively mapping RNA polyadenylation^25^. JTE-607 treatment induced significant APA changes in 921 genes, of which 847 genes (92%) shifted from a proximal PAS to a distal one (blue dots, Fig. 5a and see Methods for details). An example was shown in Fig. 5b: the proximal PAS was predominantly used for *Ptp4a1* transcripts in DMSO-treated cells. However, polyadenylation shifted to a distal PAS in JTE-607 treated cells, leading to 3′ UTR lengthening. 74 genes showed APA changes in the opposite direction (red dots, Fig. 5a), as exemplified by *Paqr8* (Fig. 5c).

**Fig 5.**
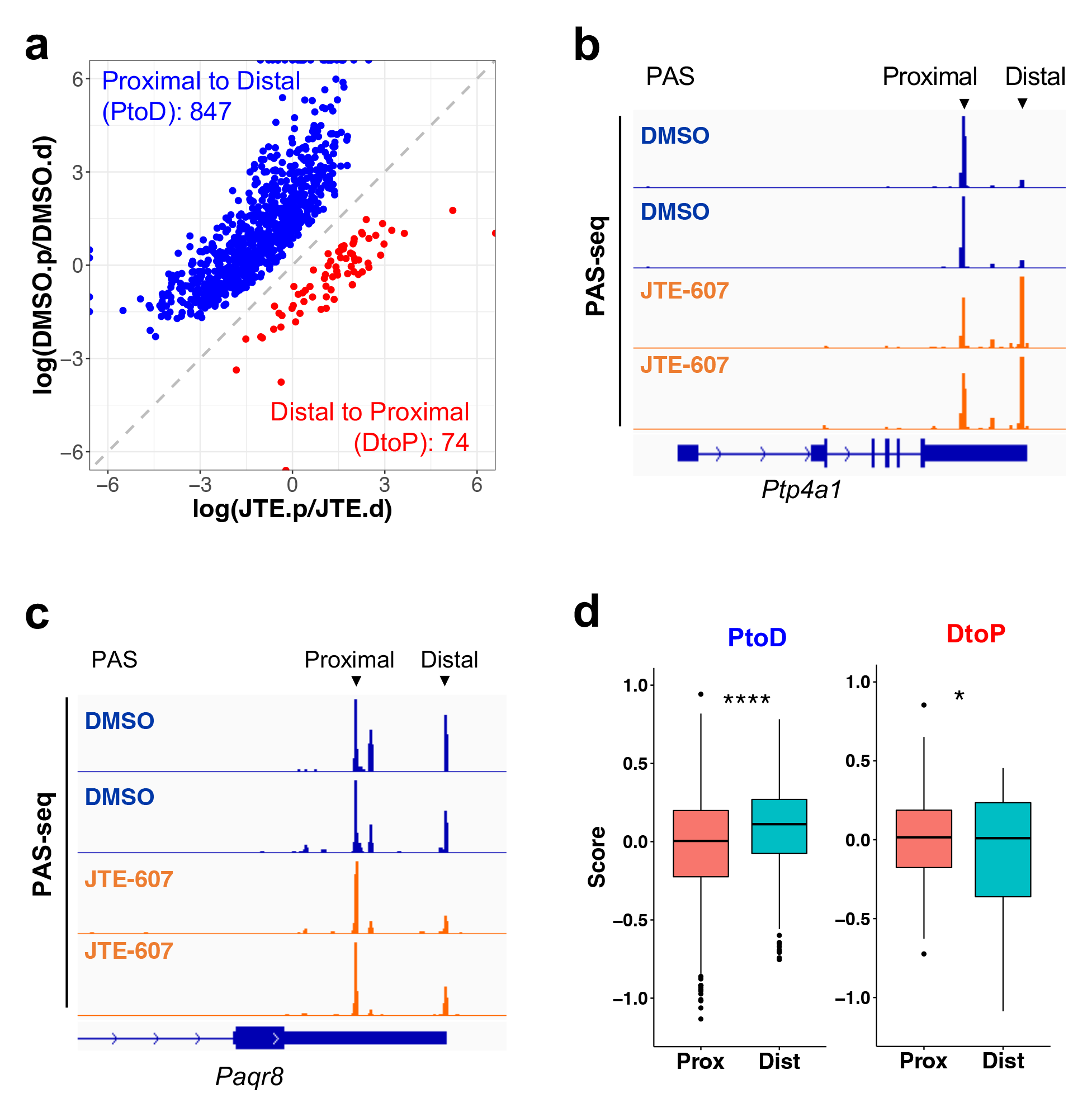
JTE-607-induced APA changes in cells are sequence-specific. **(a)** A scatter plot showing JTE-607-induced APA changes in cell. **(b-c)** PAS-seq tracks of 2 example genes: *Ptp4a1* and *Paqr*8. Two replicates for each treatment are shown and the positions of the proximal and distal PASs are marked. **(d)** Boxplots comparing the C3PO-predicted resistance scores for the proximal (Prox) and distal (Dist) PASs for the PtoD and DtoP genes. ****: p value < 0.0001; *: p value < 0.05 (t-test).

Why did JTE-607 induce the opposite APA changes in different groups of genes? Given our finding that JTE-607-mediated inhibition of mRNA 3′ processing is sequence-specific, we predicted that JTE-607 treatment would decrease the usage of the more sensitive PASs in a given gene while the usage of resistant PASs would be less impacted, leading to a net shift to the more resistant PASs. Therefore, we hypothesized that the directionality of JTE-607-induced APA change in any given gene is determined by the relative sensitivities of its alternative PASs. To test this hypothesis, we predicted the resistance scores of all annotated PASs in the human genome using C3PO and compared the scores of the proximal and distal PASs of the 921 genes that displayed significant APA shifts in JTE-607 treated cells. Interestingly, for the 847 genes that showed a shift to the distal PAS in JTE-607-treated cells, their proximal PASs are significantly more sensitive to JTE-607 than their distal ones (p < 2.2 x 10^-16^, t-test, Fig. 5d, left panel). The opposite trend was observed for the 74 genes that showed a distal-to-proximal shift (p = 0.03, t-test, Fig. 5d, right panel). Therefore JTE-607 indeed inhibited the usage of more sensitive PASs, resulting in higher usage of resistant PASs. These data confirmed that JTE-607 modulates PAS selection globally in a sequence-dependent manner in human cells and showed that JTE-607- induced APA changes depend on the relative drug sensitivities of the alternative PASs.

Additionally, we also monitored transcription termination by nascent RNA sequencing using 4-thiouridine labeled RNA (4sU-seq) in DMSO-or JTE-607-treated HepG2 cells. As mRNA 3′ processing is coupled to transcription termination, transcription termination efficiency at PAS can be used as a proxy for mRNA 3’ processing efficiency^26^. Our 4sU-seq analyses showed that JTE-607 treatment induced a global transcription termination defect (Fig. 6a). However, the levels of JTE-607-induced transcription readthrough (RT) varied widely at different PASs (Fig. 6a). For example, RT increased dramatically downstream of the PAS of the *Eif4ebp1* gene (Fig. 6b, left panel) while little change was observed for *Cox8A* gene (Fig. 6b, right panel). Thus 4sU-seq data further demonstrated that mRNA 3′ processing displayed sequence-specific sensitivity to JTE- 607-mediated inhibition in human cells.

**Fig 6.**
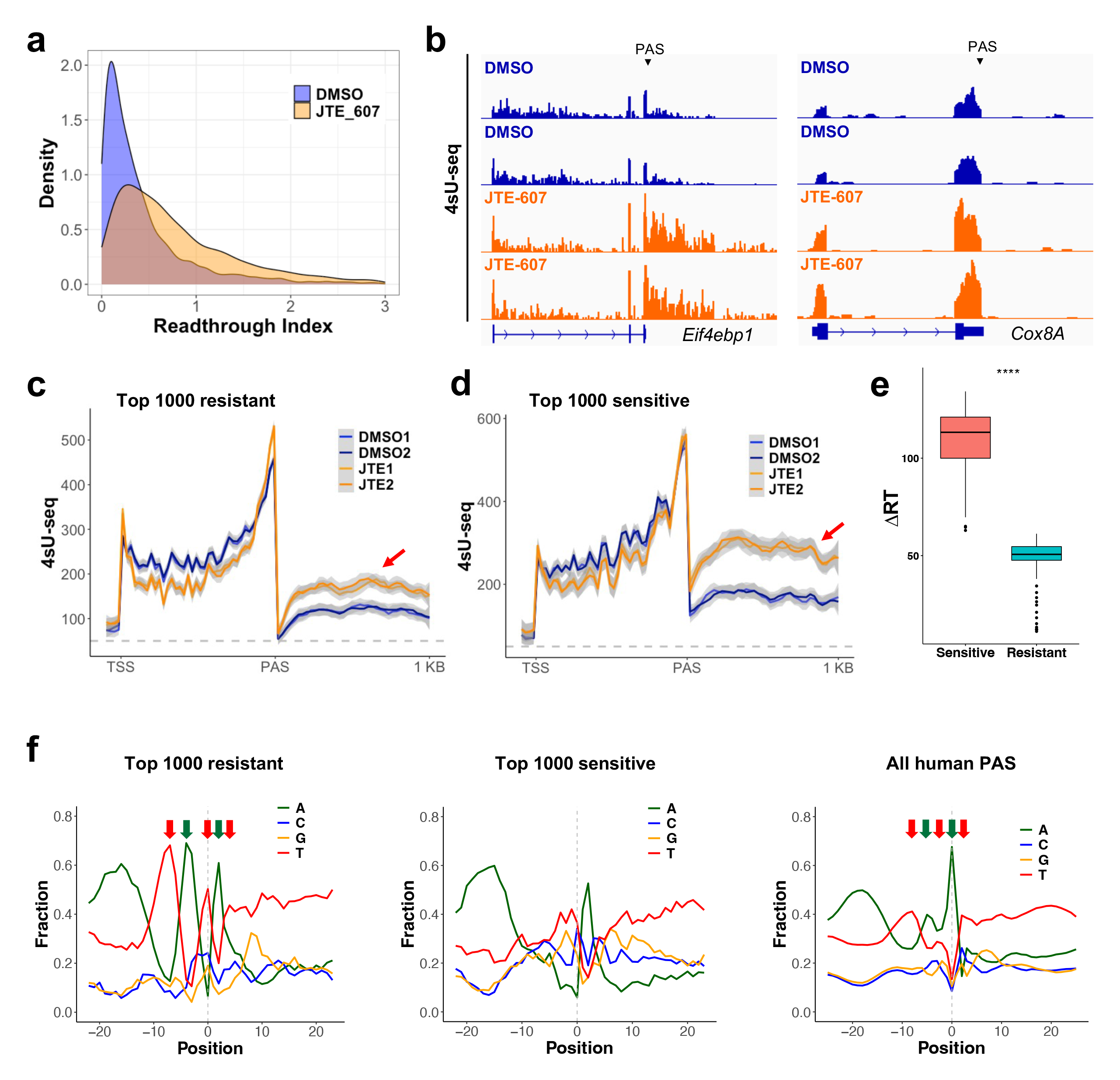
JTE-607-mediated inhibition of mRNA 3’ processing in cells is sequence-specific. **(a).** A density plot of transcription readthrough index (read counts in the 1kb downstream region/read counts in gene body) for DMSO-and JTE-607-treated cells based on 4sU-seq data. **(b).** 4sU-seq tracks for *Eif4ebp1* and *Cox8A* genes. Two replicates for DMSO and JTE-607 are shown. PAS positions are marked. **(c)** Average normalized 4sU-seq signals for the genes with the top 1,000 most resistant PASs. **(d)** Similar to C, but for the top 1,000 most sensitive PASs. Red arrow denotes region downstream of PAS. **(e)** A blox plot compare the ι1RT (the difference in 4sU-seq signals in the 1kb region downstream of the PAS. ****: p value < 0.0001, Wilcoxan test. **(f)** CS region nucleotide distribution for the top 1000 most resistant (left) and most sensitive (middle), and all human PASs (right). The T-and A-rich regions are marked by red and green arrows.

We then tested if C3PO can predict the transcription termination efficiency in JTE-607- treated cells. For comparison, we selected genes with the top 1000 resistant or sensitive PASs based on the C3PO predicted resistance scores. To avoid complications from neighboring genes, we selected genes that do not overlap with other genes in the 1kb downstream region for our analyses. The average normalized 4sU-seq signals at genes with the top 1000 resistant PASs showed that transcription terminated efficiently at these PASs in both DMSO-and JTE-607-treated cells and only modest change in RT levels was observed downstream of the PASs (Fig. 6c, red arrow), suggesting that these PASs are indeed resistant to JTE-607. By sharp contrast, for genes with the top 1000 sensitive PASs, their global 4sU-seq signals revealed significantly higher RT in JTE-607-treated cells compared to DMSO-treated cells (Fig. 6d, red arrow), suggesting that JTE- 607 induced significant inhibition of mRNA 3′ processing at these PASs. The JTE-607-induced RT levels between the sensitive and resistant PASs were highly significant (Fig. 6e, p < 2.2 x 10^-16^, Wilcoxon test). Together, our PAS-seq and 4sU-seq analyses suggest that JTE-607 inhibits mRNA 3′ processing and transcription termination in a sequence-dependent manner and that C3PO can predict the effect of JTE-607 on PAS selection and transcription termination.

Nucleotide composition of the resistant and sensitive human PASs revealed distinct patterns. JTE-607-resistant PASs have alternating U-and A-rich regions (Fig. 6f, left panel) whereas the JTE-607-sensitive PASs are generally U/G-rich (Fig. 6f, middle panel). These patterns are very consistent with the top resistant and sensitive PASs from our MPIVA screen (Fig. 3f-g). Interestingly, the average nucleotide composition of the CS regions of all annotated human PASs also displayed alternating U-and A-rich regions (Fig. 6f, right panel), suggesting that a significant portion of the human PASs are potentially resistant to JTE-607. Finally, a comparison of the resistant and sensitive PASs revealed that the resistant PASs are more conserved than the sensitive PASs (Fig. S9), indicating that the resistant PASs may be under greater selection pressure.

## Discussion

In this study, we have set out to characterize the mechanism of action for JTE-607, a novel inhibitor of the endonuclease for mRNA 3′ processing, CPSF73. Although CPSF73 is universally required for mRNA 3′ processing, we have unexpectedly discovered that Compound 2, the active form of JTE-607, inhibits the cleavage step of mRNA 3′ processing in a sequence-dependent manner both in vitro and in cells, and that the CS region sequence is a major determinant of Compound 2 sensitivity. We have comprehensively characterized the relationship between the CS region sequence and Compound 2 sensitivity using MPIVA coupled with machine learning. Our machine learning model C3PO can predict Compound 2 sensitivity based on CS sequence and the impact of JTE-607 on APA and transcription termination in human cells. Therefore, our study not only provided new insights into the mRNA 3′ processing machinery, but may also have important implications for the use of JTE-607 as a research and therapeutic tool. Furthermore, from a technological perspective, our approach described here should be broadly applicable to the studies of other small molecule modulators of gene expression.

What is the molecular mechanism for the sequence-specific sensitivity to Compound 2? Since both Compound 2 and the RNA in the CS region bind to CPSF73 at or near its active site^18,27^, these interactions are most likely mutually exclusive (Fig. 7). Thus, if a CS region RNA can bind to CPSF73 with a high affinity, it may out-compete Compound 2, rendering this PAS resistant to the drug (Fig. 7, left panel). For low-affinity CS region RNA sequences, Compound 2 bound to CPSF73 near its active site can block access by the RNA due to its low affinity, thus inhibiting cleavage (Fig. 7, right panel). Based on the structure of the histone mRNA cleavage complex^27^, which contains CPSF73 as its endonuclease, CPSF73 binds to RNA substrates via a cleft between the β-CASP domain and the metallolactamase domain. However, this cleft can only accommodate a ∼7 nt sequence, much shorter than the ∼20 nt CS region that we identified. Thus, additional mRNA 3′ processing factors likely bind to the CS region as well. Potential candidates include CPSF100 and symplekin, which form the nuclease module, or mCF, with CPSF73^7,27^. Indeed, the histone mRNA cleavage complex structure revealed that these proteins form an RNA-binding channel that can bind to ∼20 nt sequence (Fig. 7). Other mRNA 3′ processing factors can also be involved, including Fip1 and PAP. Fip1 is known to bind to U-rich sequences near the AAUAAA hexamer^28,29^. and the Compound 2-resistant CS sequences contain U-rich sequences (Fig. 3f-g). Finally, an early biochemical study showed that PAP is required for in vitro cleavage of L3 PAS, but not for SVL and that the CS region sequences determine its PAP dependency^30^. Given the important roles for the CS region in determining both PAP dependency and Compound 2 sensitivity, it is possible that PAP is involved in binding to CS region sequences. Based on these results, we propose that CPSF73 and other mRNA 3′ processing factors form an RNA-binding channel that directly binds to the CS region RNA and that this channel has sequence specificity (Fig. 7).

**Fig 7.**
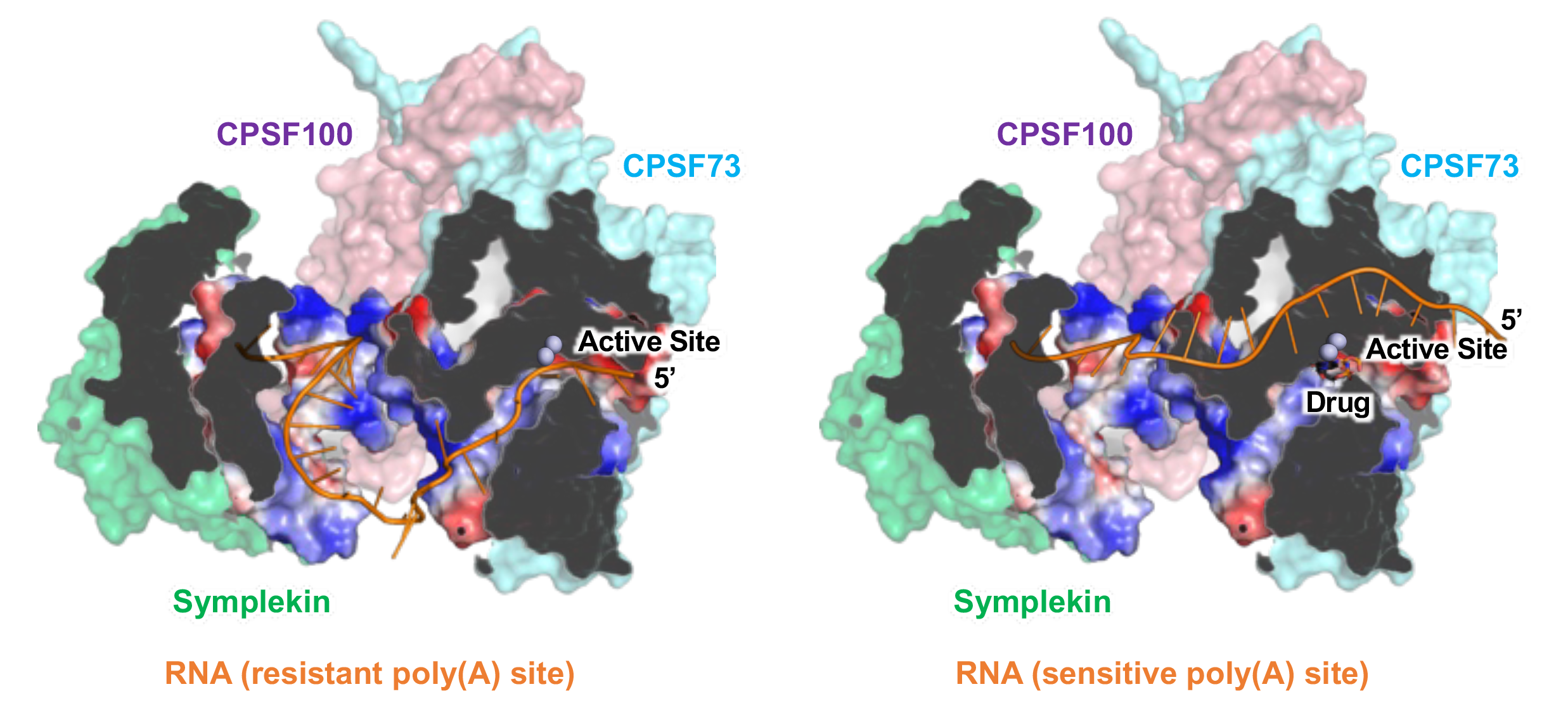
A model for sequence-specific inhibition of pre-mRNA 3’ processing by Compound 2. Artistic rendering of resistant (left panel) or sensitive (right panel) PAS RNAs within the pre-mRNA 3’ processing complex. CPSF73, CPSF100, and symplekin are shown and their structures are partially based on the histone mRNA cleavage complex (PDB accession: 6V4X), and the RNA-binding channel formed by the three proteins is highlighted. The RNA (orange thread), active site of CPSF73, and the drug (Compound 2) are marked. Please see discussion for details.

The nucleotide composition in the CS region has been conserved from yeast to human^9–11^. and this pattern is highly similar to that of the Compound 2-resistant PASs (Fig. 6f). Additionally, our data suggests that Compound 2-resistant PASs are more evolutionarily conserved than the sensitive sites (Fig. S8). It remains unclear what, if any, selection pressure can favor PASs that are resistant to a small molecule that is not present in most environments. We propose two possible models. First, Compound 2 activity may be similar to that of a chemical that is more universally found in cells. A number of small molecules, including inositol hexakisphosphate, can bind to and modulate the activities of molecular machinery in the gene expression pathway, such as the spliceosome^31^, the Integrator complex^32^, and the mRNA export factors^33^. It is possible that a naturally occurring Compound 2-like small molecule can inhibit pre-mRNA 3′ processing and many PASs evolved to overcome such inhibition. Secondly, the CS region sequence may impact transcription termination independently of its effect on cleavage efficiency. According to our model, the resistant PASs interact with CPSF73 and other mRNA 3′ processing factors more strongly (Fig. 7). Because the mRNA 3′ processing machinery is known to directly bind to RNA polymerase II^34–36^, such interaction could contribute to slowing down the polymerase, thus promoting termination. Thus, a subset of PASs may have evolved to stimulate transcription termination and the Compound 2 resistance is an unintended consequence of such evolution.

In addition to CS region sequence, we have provided evidence that UPS sequence as well as RNA secondary structure may also contribute to Compound 2 sensitivity, albeit in a context-dependent manner (Fig. 2a and Fig. S8). UPS sequence can modulate CPSF73-CS region RNA interactions indirectly through associated protein factors. Alternatively, UPS could form secondary structures with the CS region, thus impacting its interactions with CPSF73 more directly. In fact, the mCF module has been shown to bind directly to double-stranded RNAs in the histone mRNA cleavage complex^27^. Additionally, secondary structures are widespread in PAS regions and have been shown to modulate mRNA 3′ processing.^23^ Thus, it is possible that RNA secondary structures within a PAS could impact not only its cleavage efficiency, but also its Compound 2 sensitivity. Further studies are needed to fully elucidate the roles of UPS and RNA secondary structures in drug sensitivity.

Our results may have implications for understanding how JTE-607 specifically kills myeloid leukemia and Ewing’s sarcoma cell lines. As mentioned earlier, JTE-607 is a pro-drug and is converted to Compound 2 by the cellular enzyme CES1^18^. Although cellular CES1 levels may contribute to the cell type specificity, previous studies showed that CES1 level is a poor predictor for JTE-607 sensitivity^18^. Thus, the molecular basis for cell type-specific toxicity of JTE- 607 remains unknown. Based on the results reported here, we propose two possible mechanisms for explaining the cell type-specific drug sensitivity. First, the potency for JTE-607-mediated inhibition of mRNA 3′ processing may be cell type-specific. Our model suggests that the drug sensitivity is determined by the interaction affinity between the CPSF73 and other mRNA 3′ processing factors and the CS region sequence. If cell type-specific mechanisms can modulate the specificity of this interaction, they can alter JTE-607 sensitivity globally. This could result from cell type-specific expression levels or post-translational modification of CPSF73 and other mRNA 3′ processing factors that bind to the CS region. For example, Fip1 levels are known to change during differentiation^28,37^. Symplekin, CPSF100, and PAP are known to be sumoylated and/or phosphorylated^38–40^. It will be important to determine if these factors display different expression levels or post-translational modifications between JTE-607-sensitive and -resistant cell types. Alternatively, the sequence specificity of JTE-607 is similar among different cell types. However, myeloid leukemia and Ewing’s sarcoma cells may be uniquely dependent on one gene or a subset of genes whose PASs are highly sensitive to JTE-607. For example, a recent study identified PDXK, an enzyme in the vitamin B6 metabolism pathway, as a unique acute myeloid leukemia dependency gene^41^. If the PASs of such dependency genes are sensitive to JTE-607, the expression of these genes would be repressed by JTE-607 treatment, leading to cell death in specific cell types. Further studies are needed to distinguish between these models and the results will have significant implications on how to improve the efficacy of this compound as a potential anti-cancer and anti-inflammation therapy.

## Acknowledgement

We would like to thank Dr. Rohan Beckwith for providing reagents. We wish to acknowledge the support of the Chao Family Comprehensive Cancer Center Shared Resource Genomics High-Throughput Facility, supported by the National Cancer Institute of the National Institutes of Health under award number P30CA062203. This study was supported by the following grants: NIH AI166703 and GM090056. L.L. is supported by the Center for Virus Research Graduate Fellowship provided by the UCI Division of Graduate Studies. A.MY. is a Washington Research Foundation Postdoctoral Fellow

## Methods

### Cell Culture and JTE-607 Treatment Condition

HepG2 cell line were cultured in Dulbecco’s modified Eagle medium (DMEM) supplemented with 10% (v/v) fetal bovine serum (FBS). Cells were incubated at 37 °C in a 5% (v/v) CO_2_-enriched incubator. Suspension HeLa S3 cell (a kind gift from Dr. Bert Semler, UC Irvine) was maintained in Joklik Modified MEM (JEME) supplemented with 2.4 mM sodium bicarbonate and 8% (v/v) newborn calf serum (NCS) in a spinner flask at 37°C with ambient CO_2_. For JTE-607 treatment, 20 µM final concentration of JTE-607 (Tocris) in neat DMSO was added to the cell culture media and incubated at 37°C for 4 hours.

### Large-scale HeLa nuclear extract (NE)

Large-scale HeLa nuclear extract (NE) was made as previously described (Abmayr et al., 2006) with minor modifications. Briefly, 10 liters of spinner HeLa cells were pelleted by centrifugation. The cells were swelled on ice using hypotonic buffer A (10 mM HEPES-NaOH pH 7.9, 10 mM KCl, 1.5 mM MgCl_2_, 10 mM 2-Mercaptoethanol) and then dounce homogenized with 15 strokes using a type B pestle. Each stroke involves a 30-second motion containing one 15-second up and one 15-second down motion. Cell lysis was closely monitored by mixing a small aliquot of cells with trypan blue and observing under light microscope. Dounce homogenization was stopped when ∼85% cell lysis was achieved. Nuclei were pelleted, extracted with high salt buffer C (20 mM HEPES-NaOH pH 7.9, 420 mM NaCl, 1.5 mM MgCl_2_, 0.2 mM EDTA, 25% glycerol, 10 mM 2-Mercaptoethanol, 0.5 mM PMSF) freshly supplemented with 1X Halt proteinase inhibitor cocktail (Thermo) at 4°C for one hour with constant rotation. The extracted nuclei were pelleted, and the supernatant (NE) was dialyzed twice against 60 volume of buffer D100 (20 mM HEPES-NaOH pH 7.9, 100 mM KCl, 1 mM MgCl_2_, 0.2 mM EDTA, 10% glycerol 10 mM 2- Mercaptoethanol, 0.5 mM PMSF) at 4°C for 1.5 hours each time. After dialysis, the NE was aliquoted, flash frozen on dry ice and stored at −80°C until use.

### In vitro Cleavage Assay

All PASs were cloned into the pBlueScript II KS+ vector. RNA substrates were synthesized by run off in vitro transcription (IVT) using T7 polymerase (NEB) in the presence of [α-^32^P]-UTP according to the manufacture’s protocol. For in vitro cleavage reaction with Compound 2, the NE was pre-incubated with 10% DMSO or various concentration of Compound 2 (0.1, 0.5, 2.5, 12.5, 62.5, 100 µM) in 10% DMSO for 30 minutes on ice before the other components were added. Each in vitro cleavage reaction is a 10µl reaction containing 20 cps radiolabeled pre-mRNA, 44% (v/v) HeLa NE, 8.8 mM HEPES-OH (pH 7.9), 44 mM KCl, 0.44 mM MgCl_2_, 0.2 mM 3′-dATP (Sigma), 2.5% (v/v) polyvinyl alcohol (PVA), 40 mM creatine phosphate, 4 mM 2- Mercaptoethanol, and 1% (v/v) DMSO or Compound 2. Cleavage was carried out for 90 minutes at 30°C. Proteinase K digestion mix (30 mM Tris-HCl pH 7.9, 10 mM EDTA, 1% SDS, 0.1 µg/µl proteinase K, 0.05 µg/µl yeast tRNA) was then added to halt the reaction and the samples were incubated at 37°C for 15 min. RNA was then phenol chloroform extracted and resolved on an 8% Urea-PAGE at 800 V for 45 minuets in TBE. Gel was then transferred to a filter paper, dried at 80°C for 30 minutes, exposed to a phosphoscreen overnight and visualized by phosphorimaging. IC_50_ was calculated using the equation: [Inhibitor] vs. normalized response -- Variable slope on Prism.

We have found that this assay is very sensitive to the strength of RNA radioactivity and freshness of NE. It is recommended that freshly purchased [α-^32^P]-UTP (less than a week old) and NE made and stored at −80°C for fewer than two months to be used for this assay.

### Electrophoretic mobility shift assay (EMSA)

NE was pre-incubated with DMSO or Compound 2 as described in the in vitro cleavage assay. Gel shift is performed in a 10µl reaction containing 20 cps radiolabeled RNA, 1 mM ATP, 20 mM creatine phosphate, 10 µg/µl yeast tRNA, 44% HeLa NE, and 1% (v/v) DMSO or Compound 2. The reaction mixture was incubated for 20 minutes at 30°C and immediately cooled on ice for 2 minutes. Heparin was added to 0.4 µg/µl and the reaction was incubated for an additional 5 minutes on ice. 5µl of the reaction was resolved on 4% native PAGE in 1x Tris-Glycine running buffer (pH 8.3) at 100V for 4 hours in an ice bath. Gel was dried and visualized the same as in vitro cleavage assay described above.

### Massively Parallel in vitro Assay (MPIVA)

#### Cloning

L3 and SVL containing 23 random nucleotides CS spanning YA cleavage position was purchased from IDT as ssDNA oligo and PCR amplified to generate dsDNA. The dsDNA library was cloned into pBlueScript II KS+ vector by Gibson Assembly (NEB) and electroporated into ElectroMAX DH5α (Thermo). Plasmid library size, structure, and diversity were determined as previously described^42^.

#### Coupled in vitro Cleavage and Polyadenylation Assay

RNA libraries were synthesized by run off IVT using T7 polymerase (NEB) according to the manufacture’s protocol followed by treatment with RQ1 DNase (Promega) to remove DNA template. The RNA pool was purified by phenol chloroform extraction and was either polyadenylated (for input) or 3′-dATP blocked (for DMSO and Compound 2 treated) by *E. coli* PAP (NEB). The RNAs were then undergo a coupled cleavage and polyadenylation assay in multiple 600µl reactions containing 6 pmol RNA, 44% (v/v) HeLa NE, 8.8 mM HEPES-OH (pH 7.9), 44 mM KCl, 1.44 mM MgCl_2_, 1 mM ATP, 2.5% (v/v) polyvinyl alcohol (PVA), 20 mM creatine phosphate, 4 mM 2-Mercaptoethanol, and either 1% DMSO or 0.5 µM, 2.5 µM, 12.5 µM Compound 2 in DMSO. The reaction mixture was incubated for 90 min at 30°C, proteinase K digested as described above for regular in vitro cleavage assay, except that proteinase K was raised to 3µg/µl, and then phenol chloroform extracted.

#### MPIVA Sequencing Library Construction

The phenol chloroform extracted RNA from previous step weas further purified to select for polyadenylated RNA using NEBNext Poly(A) mRNA Magnetic Isolation Module (NEB) and reverse transcribed using SuperScript III reverse transcriptase (Invitrogen) with an anchored oligo dT primer. Library cDNA was beads purified (Beckman Coulter) and amplified using a library-specific forward primer and reverse primer containing Illumina adaptor sequences and a region that matches part of the sequence added during RT. The amplified libraries were resolved on a 2.5% low melting point agarose gel and extracted.

#### MPIVA RNA-seq read alignment

All MPIVA read 1 and read 2 FASTQ files were merged using bbmerge V38 using option ` maxloose=t`^43^. Untreated RNA-seq reads were used to establish the sequences of the full randomized PAS region contained in the IVT pool. All sequences of the 25 nt randomized region were clustered using starcode version 1.4^44^ to account for sequencing errors and determine consensus sequences of this randomized region. The next steps are to enable assignment of expected cleaved RNAs to a unique 25 nt randomized region. The expected cleaved lengths of the 25 nt consensus sequences for L3 and SVL backbones (13 nt and 12 nt, respectively) were used as unique identifiers of the full randomized region. If any of these identifiers were not unique within L3 and SVL libraries, respectively, then these sequences were not used in subsequent analyses.

Next, RNA-seq reads from DMSO and drug-treated libraries were locally aligned against the shared 5′ region of the reporter constructs to determine the beginning of the randomized region. The part of the RNA-seq read containing the randomized region and shared 3′ region was locally aligned against the list of consensus sequences. Only reads with a unique alignment to a single consensus sequence were kept, and cut sites were also determined from this alignment. Additional checks were performed to ensure cut sites are not misassigned inside the poly(A) tail due to an adenine in the reference and these cases were corrected if found. Sequences with at least 50 reads in the DMSO libraries were kept to avoid noise from lowly abundant RNA sequences in the IVT pool. 158,298 L3 variants and 103,018 SVL variants were left after this read depth filtering step. A pseudocount of 1 was then added to variants in L3 2.5 μM, L3 12.5 μM, and SVL 12.5 μM due to drug-mediated drop out of high abundance variants in the DMSO libraries. This pseudocount avoids having undefined drug sensitivities in later steps that would be introduced by log(0). Each variant that passed these checks were counted and converted to a percentage within each RNA-seq library to account for sequencing depth by dividing by the total number of kept reads.

Drug sensitivity for each variant in each dose of Compound 2 was defined as the log ratio of normalized reads from drug-treated RNA-seq divided by the normalized reads from DMSO-treated RNA-seq. Within a given drug dose, sequences with higher log ratios are more resistant than those sequences with lower log ratios.

#### Minimum free energy folding of IVT RNA’s

Minimum free energy (MFE) predictions were done with RNAStructure version 6.4’s *Fold*^45^ and the entire IVT RNA sequence was used. ΔG of each MFE were determined with RNAStructure’s *efn2* with command line argument `--simplè. ΔG’s from the top 10,000 resistant and sensitive sequences were compared (Quantification and statistical analysis).

#### C3PO machine learning architecture and training

The architecture chosen for predicting drug sensitivity is based on a previously published 3-layer convolutional neural network (CNN) designed for predicting polysome profiles.^22^ The model takes in 25 nt one-hot encoded sequences followed by:

First convolution layer: 120 filters (8 × 4), batch normalization, ReLU activation, zero-padding to maintain the same length input and output, and 0% dropout.

Second convolution layer: 120 filters (8 × 1), batch normalization, ReLU activation, zero-padding to maintain the same length input and output, and 0% dropout.

Third convolution layer: 120 filters (8 × 1), batch normalization, ReLU activation, zero-padding to maintain the same length input and output, and 0% dropout.

Dense layer: 80 nodes, batch normalization, ReLU activation and 10% dropout. Output layer: 3 linear outputs.

The Adam optimizer^46^ was used for model fitting with a mean squared error loss function, batch size of 64, and sample weights based on DMSO read depth.

Sequences were assembled into test and training sets to mix highly covered variants from both RNA contexts (L3 and SVL) into the test and training sets. Within each RNA context, sequences were ordered by DMSO read depth, then split based on the sequences’ number in this ordering into odd and even lists, and then the odd and even lists were concatenated together. This odd/even splitting is to include high coverage sequences in both the training and test sets. Finally, the L3 and SVL sequences were interleaved to make an even coverage between RNA contexts in the test set. The top 4,120 sequences were used as the test set with the remaining sequences used as the training set. The 4,120 test set size was chosen because it reflects 2% of the variant space in SVL which contains less variants than L3.

Ten iterations of training with 6 epochs were conducted to account for slight variations in model performance due to stochasticity in the training algorithm. Performance between iterations were evaluated by the square of Pearson’s r (R^2^) between measured and predicted Compound 2 sensitivity in the test sequences. The best performing iteration was kept and used in further analyses.

#### Exploring additional machine learning architectures and training

Additional deep learning and training pipelines were explored based on CNN’s and Dilated Residual Networks. With the C3PO CNN architecture, training was done with 4-8 epochs and these number of epochs performed relatively similarly on the test set (SI Models Table). We also explored using a validation set (4,120 sequences) derived from the training set to determine an early stopping criterion for the number of epochs trained, and this performed similarly to the models trained with a preset number of epochs. Among the three drug doses predictions, 12.5 μM predictions performed better leading us to train models for only this dose as well. However, 12.5 μM prediction performance between the three dose predictions and the one dose prediction were negligibly different so we used the model with three dose predictions.

Hyperband training^47^ with the CNN architecture was also performed to ascertain potential optimal hyperparameter values. Hyperparameters were allowed to range from 1-5 1D convolutional layers with ReLU activation and batch normalization; 8-140 (step 16) number of filters; followed by pooling choices of average, max, or none; and dropout rates of 0-0.5 (step 0.1). These convolutional layer(s) are followed by a Flatten layer, and 1-3 dense layers. Each dense layer can be of size 20-200 (step 20) with ReLU activation, batch normalization, and dropout rates of 0-0.5 (step 0.1). Learning rate parameters were also allowed to range between 1×10^-5^-1×10^-1^. Training was allowed to stop early based on the validation set’s mean squared error and a minimum delta of 0.001 and patience of 5 epochs. Hyperband training was done with an output layer for all three drug doses, and hyperband training was also tried with an output layer for only predicting12.5 μM Compound 2 resistance.

Due to the improvement of APARENT2^24^ which is a residual neural network (ResNet) over APARENT^7^ which is a CNN, we also tried an architecture similar to APARENT2 with our task on predicting Compound 2 sensitivity. We tested the residual neural network architecture with predicting both Compound 2 sensitivity and cleavage patterns with the hypothesis that learning sequence features that affect cleavage site usage would improve the Compound 2 sensitivity predictions. Input to the residual network is a one-hot encoded 25 nt sequence which is the same as our CNN models and is followed by 20 residual blocks where each block contains 2 layers of dilated convolutions and a skip connection. More specifically, there are 5 residual groups where each residual group contains 4 residual blocks with 32 channels and convolutional filters of size 3. Each residual block is encoded the same as APARENT2 where each blocks has two one-dimensional convolutional layers with batch-normalization, ReLU activation, and a filter dilation rate. There are additional skip connections from between each residual group to the last convolutional layer and produces a vector of length 26, s(x). The 26^th^ position is for all cuts found at positions not found the 25 nt randomized region. For training and accounting for any background sequence biases, a boolean is passed to indicate whether the data point is from the L3 or SVL background which is multiplied with a position-specific weight matrix and linearly combined with s(x). We also kept APARENT2’s random shifting of the input sequence and cleavage distribution during training to force the network to not simply learn the designed expected cleavage position in each library. These scores containing library-specific information are sent to four different linear dense layers for separate predictions of cleavage profiles of all four drug doses and softmax transformation is applied to each. For Compound 2 sensitivity prediction, s(x) undergoes average pooling, and the library indicator is concatenated before a linear dense layer for final output. KL-divergence is used as the loss function for cleavage profiles and mean squared error for Compound 2 sensitivities. Total loss is a weighted average of half from Compound 2 sensitivities, and the other half split evenly between the four cleavage profiles. The ResNet was trained with Keras’s implementation of the Adam optimizer, batch size of 64, stopping criteria based on a validation set (4,120 sequences) derived from the training set.

We first tried 1, 2, 4, 2, and 1 as dilation rates for the 5 residual groups and performed similarly to previously trained CNNs with R^2^ values of 0.232, 0.541, and. 0.681 for the three Compound 2 doses but did not outperform C3PO (SI Table 1). We also tried lower dilation rates of 1, 2, 2, 2, and 1 as well as 1, 1, 1, 1, and 1 which performed worse. Using the dilation rates 1, 2, 4, 2, and 1, we trained for exactly 7 epochs and did not find improved performance. We also increased the cleavage profile length to 27 to separately model cuts found at positions greater than the 25 nt randomized region in position 26, and position 27 is filled when a sequence is not found at a given Compound 2 dose (i.e. sensitive sequences that drop out at higher Compound 2 doses). This led to R^2^ values of 0.229, 0.55, and. 0.686 for the three Compound 2 doses which also did not outperform C3PO (SI Table 1). Finally, we increased the weight of Compound 2 sensitivity predictions to 75% of the total loss which did not lead to better performance than C3PO.

#### Convolutional layers 1 and 2 activation analysis

Convolutional layers 1 and 2 were analyzed similarly to a previously published analysis of a CNN that predicts alternative polyadenylation (APARENT)^42^. In brief, every filter in both convolutional layers were correlated with predictions of drug sensitivity at the 12.5 μM dose. The top 5,000 input sequences from the training set that achieved maximal filter activation were put into a position weight matrix and used to generate position-aware consensus sequence logos^48^. Pearson’s r plots of each filter’s activations with predicted 12.5 μM Compound 2 sensitivity at each position are plotted below these filter-specific sequence logos. Layer 1 filters are 8 positions wide, and layer 2 filters are 15 positions wide. Note that the convolutional layers in C3PO contain even zero-padding to maintain an input/output size of 25. The padding should be accounted for when analyzing the filters’ Pearson r plots. For example in layer 1, the sequences are padded with 4 0’s on both the left and right.

#### APARENT2 predictions and comparisons

APARENT2 predictions of logodds of cleavage at expected cleavage position versus elsewhere were done on all MPRA sequences, centered at their expected cut site which is the expected format of APARENT2. Predictions with read depth of at least 150 in the Input libraries were kept for further analysis. APARENT2 predictions were compared against the logodds of expected cleaved DMSO read counts and Input read counts which estimates the *in vitro* cleavage efficiency. Additionally, APARENT2 predictions were compared against the logodds of expected cleaved 12.5 μM Compound 2 read counts and Input read counts which estimates the *in vitro* drug resistance.

### 4sU-seq

HepG2 cells were treated with DMSO or 20 µM JTE-607 (Tocris) for 3 hours at 37°C. 500 µM 4sU (Sigma) was then added to the DMSO/JTE-607 containing media and cells were incubated at 37°C for one additional hour. After incubation, cells were lysed in Trizol (Invitrogen), and total RNA was extracted following the manufacturer’s protocol. 4sU RNA enrichment and library preparation were done as previously described with minor modifications^49^. Briefly, 50 µg total RNA was used as the starting material and biotinylated with biotin-HPDP (Thermo). 4sU labeled and biotinylated RNA was enriched with streptavidin beads by rotating at room temperature for 1.5 hours, eluted with 100 mM DTT, and further purified by phenol chloroform extraction.

### PAS-seq

HepG2 cells were treated with DMSO or 20 µM JTE-607 (Tocris) for 4 hours at 37°C and total RNA was extracted by Trizol (Invitrogen). 10 µg of total RNA was used to prepare PAS-seq libraries as previously described^50^. Briefly, 10 µg total RNA was fragmented using fragmentation buffer (Thermo) and reverse transcribed by SuperScript III (Thermo). The cDNA was circularized by Circligase (Lucigen) and then re-linearized by BamHI. The digested DNA was then PCR amplified and a ∼200 bp region was gel extracted and sequenced. During the execution of this study, we have further optimized the library preparation steps of the protocol and for the most updated PAS-seq protocol please refer to PAS-seq 2^51^.

### 4sU-seq and PAS-seq data analysis

For 4sU-seq, reads were mapped to human hg19 using STAR^52^ and bigwig files were generated using deepTools^53^. For PAS-seq, reads without a poly(A) tail (fewer than 15 consecutive A’s) were removed. The polyA tail sequence and linker sequence was trimmed from remaining reads before mapping. The trimmed reads were mapped to human hg19 using STAR^52^ as done for 4sU-seq except that the EndToEnd parameter was used only for PAS-seq. The resulting bam output file was converted to a bed file using BEDTools^54^. Reads that may have been due to internal priming (reads where there were 6 consecutive A’s within 10 nucleotides downstream of the PAS, or 7 A’s out of 10 nucleotides downstream of the PAS) were removed. The resulting bed file was then converted back to a bam file using BEDTools.^54^ The location of the 3′ end of each read was extracted using BEDTools^54^ and was then compared to the location of all annotated PAS within PolyA_DB^55^ to retrieve read counts for each PAS. APA analysis was performed by edgeR^56^.

### Quantification and statistical analysis

Pearson’s r and R^2^ (square of Pearson’s r) are used in Figures 1∼4, S2, S3, S6 and S7 and related text as well as SI Models Table. Potential inequality of the top 10,000 resistant and sensitive sequences’ MFE ΔG’s were tested with a two-sided t-test with unequal variance. 6-mers in the top 10,000 resistant and sensitive sequences were found to be significant by a binomial test with a null hypothesis of probability of success = 0.25^6^ and alternative hypothesis of > 0.25^6^. p-value threshold was adjusted by the number of possible k-mers, 4^6^, and thus significant 6-mers must have p-values 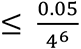.

## Reference

1. Colgan, D. F. & Manley, J. L. Mechanism and regulation of mRNA polyadenylation. Genes Dev 11, 2755–2766 (1997).

2. Chan, S., Choi, E. A. & Shi, Y. Pre-mRNA 3’-end processing complex assembly and function. Wiley Interdiscip Rev RNA 2, 321–335 (2011).

3. Shi, Y. Alternative polyadenylation: new insights from global analyses. RNA 18, 2105–2117 (2012).

4. Mitschka, S. & Mayr, C. Context-specific regulation and function of mRNA alternative polyadenylation. Nat. Rev. Mol. Cell Biol. (2022) doi:10.1038/s41580-022-00507-5.

5. Tian, B. & Manley, J. L. Alternative polyadenylation of mRNA precursors. Nat. Rev. Mol. Cell Biol. 18, 18–30 (2016).

6. Mandel, C. R. et al. Polyadenylation factor CPSF-73 is the pre-mRNA 3′-end-processing endonuclease. Nature 444, 953–956 (2006).

7. Shi, Y. & Manley, J. L. J. L. The end of the message: multiple protein--RNA interactions define the mRNA polyadenylation site. Genes Dev. 29, 889–897 (2015).

8. Sheets, M. D., Ogg, S. C. & Wickens, M. P. Point mutations in AAUAAA and the poly (A) addition site: effects on the accuracy and efficiency of cleavage and polyadenylation in vitro. Nucleic Acids Res 18, 5799–5805 (1990).

9. Ozsolak, F. et al. Comprehensive polyadenylation site maps in yeast and human reveal pervasive alternative polyadenylation. Cell 143, 1018–1029 (2010).

10. Liu, X. et al. Comparative analysis of alternative polyadenylation in S. cerevisiae and S. pombe. Genome Res. 27, 1685–1695 (2017).

11. Derti, A. et al. A quantitative atlas of polyadenylation in five mammals. Genome Res 22, 1173–1183 (2012).

12. Palencia, A. et al. Targeting *Toxoplasma gondii* CPSF3 as a new approach to control toxoplasmosis. EMBO Mol. Med. 9, 385–394 (2017).

13. Begolo, D. et al. The trypanocidal benzoxaborole AN7973 inhibits trypanosome mRNA processing. PLoS Pathog. 14, (2018).

14. Sonoiki, E. et al. A potent antimalarial benzoxaborole targets a Plasmodium falciparum cleavage and polyadenylation specificity factor homologue. Nat. Commun. 8, 1–11 (2017).

15. Sasaki, J. et al. Prior burn insult induces lethal acute lung injury in endotoxemic mice: Effects of cytokine inhibition. Am. J. Physiol. - Lung Cell. Mol. Physiol. 284, (2003).

16. Uesato, N., Fukui, K., Maruhashi, J., Tojo, A. & Tajima, N. JTE-607, a multiple cytokine production inhibitor, ameliorates disease in a SCID mouse xenograft acute myeloid leukemia model. Exp. Hematol. 34, 1385–1392 (2006).

17. Jian, M. Y., Koizumi, T., Tsushima, K. & Kubo, K. JTE-607, a cytokine release blocker, attenuates acid aspiration-induced lung injury in rats. Eur. J. Pharmacol. 488, 231–238 (2004).

18. Ross, N. T. et al. CPSF3-dependent pre-mRNA processing as a druggable node in AML and Ewing’s sarcoma. Nat. Chem. Biol. 16, 50–59 (2020).

19. Kakegawa, J., Sakane, N., Suzuki, K. & Yoshida, T. JTE-607, a multiple cytokine production inhibitor, targets CPSF3 and inhibits pre-mRNA processing. Biochem. Biophys. Res. Commun. 518, 32–37 (2019).

20. Boreikaite, V., Elliott, T. S., Chin, J. W. & Passmore, L. A. RBBP6 activates the pre-mRNA 3’ end processing machinery in humans. Genes Dev. 36, 210–224 (2022).

21. Gutierrez, P. A., Baughman, K., Sun, Y. & Tong, L. A real-time fluorescence assay for CPSF73, the nuclease for pre-mRNA 3’-end processing. RNA N. Y. N 27, 1148–1154 (2021).

22. Sample, P. J. et al. Human 5′ UTR design and variant effect prediction from a massively parallel translation assay. Nat. Biotechnol. 37, 803–809 (2019).

23. Wu, X. & Bartel, D. P. Widespread Influence of 3′-End Structures on Mammalian mRNA Processing and Stability. Cell 169, 905–917.e11 (2017).

24. Linder, J., Koplik, S. E., Kundaje, A. & Seelig, G. Deciphering the impact of genetic variation on human polyadenylation using APARENT2. Genome Biol. 23, 232 (2022).

25. Yoon, Y., Soles, L. V. & Shi, Y. PAS-seq 2: A fast and sensitive method for global profiling of polyadenylated RNAs. Methods Enzymol. 655, 25–35 (2021).

26. Nojima, T. et al. Mammalian NET-seq reveals genome-wide nascent transcription coupled to RNA processing. Cell 161, 526–540 (2015).

27. Sun, Y. et al. Structure of an active human histone pre-mRNA 3′-end processing machinery. Science 367, 700–703 (2020).

28. Lackford, B. et al. Fip1 regulates mRNA alternative polyadenylation to promote stem cell self-renewal. EMBO J 33, 878–889 (2014).

29. Martin, G., Gruber, A. R., Keller, W. & Zavolan, M. Genome-wide Analysis of Pre-mRNA 3’ End Processing Reveals a Decisive Role of Human Cleavage Factor I in the Regulation of 3’ UTR Length. Cell Rep. 1, 753–763 (2012).

30. Ryner, L. C., Takagaki, Y. & Manley, J. L. Sequences downstream of AAUAAA signals affect pre-mRNA cleavage and polyadenylation in vitro both directly and indirectly. Mol Cell Biol 9, 1759–1771 (1989).

31. Wan, R., Bai, R., Yan, C., Lei, J. & Shi, Y. Structures of the Catalytically Activated Yeast Spliceosome Reveal the Mechanism of Branching. Cell 177, 339–351.e13 (2019).

32. Lin, M.-H. et al. Inositol hexakisphosphate is required for Integrator function. Nat. Commun. 13, 5742 (2022).

33. York, J. D., Odom, A. R., Murphy, R., Ives, E. B. & Wente, S. R. A phospholipase C-dependent inositol polyphosphate kinase pathway required for efficient messenger RNA export. Science 285, 96–100 (1999).

34. Richard, P. & Manley, J. L. Transcription termination by nuclear RNA polymerases. Genes Dev 23, 1247–1269 (2009).

35. Bentley, D. L. Rules of engagement: co-transcriptional recruitment of pre-mRNA processing factors. Curr Opin Cell Biol 17, 251–256 (2005).

36. Proudfoot, N. J. Transcriptional termination in mammals: Stopping the RNA polymerase II juggernaut. Science 352, aad9926–aad9926 (2016).

37. Schwich, O. D. et al. SRSF3 and SRSF7 modulate 3’UTR length through suppression or activation of proximal polyadenylation sites and regulation of CFIm levels. Genome Biol. 22, (2021).

38. Vethantham, V., Rao, N. & Manley, J. L. Sumoylation regulates multiple aspects of mammalian poly(A) polymerase function. Genes Dev 22, 499–511 (2008).

39. Vethantham, V., Rao, N. & Manley, J. L. Sumoylation modulates the assembly and activity of the pre-mRNA 3’ processing complex. Mol Cell Biol 27, 8848–8858 (2007).

40. Colgan, D. F., Murthy, K. G., Prives, C. & Manley, J. L. Cell-cycle related regulation of poly(A) polymerase by phosphorylation. Nature 384, 282–285 (1996).

41. Chen, C.-C. et al. Vitamin B6 Addiction in Acute Myeloid Leukemia. Cancer Cell 37, 71–84.e7 (2020).

42. Bogard, N., Linder, J., Rosenberg, A. B. & Seelig, G. A Deep Neural Network for Predicting and Engineering Alternative Polyadenylation. Cell 178, 91-106.e23 (2019).

43. Bushnell, B., Rood, J. & Singer, E. BBMerge – Accurate paired shotgun read merging via overlap. PLOS ONE 12, e0185056 (2017).

44. Zorita, E., Cuscó, P. & Filion, G. J. Starcode: sequence clustering based on all-pairs search. Bioinforma. Oxf. Engl. 31, 1913–1919 (2015).

45. Reuter, J. S. & Mathews, D. H. RNAstructure: software for RNA secondary structure prediction and analysis. BMC Bioinformatics 11, 129 (2010).

46. Kingma, D. P. & Ba, J. Adam: A Method for Stochastic Optimization. http://arxiv.org/abs/1412.6980 (2017) doi:10.48550/arXiv.1412.6980.

47. Li, L., Jamieson, K., DeSalvo, G., Rostamizadeh, A. & Talwalkar, A. Hyperband: a novel bandit-based approach to hyperparameter optimization. J. Mach. Learn. Res. 18, 6765–6816 (2017).

48. Alipanahi, B., Delong, A., Weirauch, M. T. & Frey, B. J. Predicting the sequence specificities of DNA-and RNA-binding proteins by deep learning. Nat. Biotechnol. 33, 831– 838 (2015).

49. Wang, X. et al. Herpes simplex virus blocks host transcription termination via the bimodal activities of ICP27. Nat. Commun. 11, 293 (2020).

50. Wang, X. et al. Mechanism and consequences of herpes simplex virus 1-mediated regulation of host mRNA alternative polyadenylation. PLOS Genet. 17, e1009263 (2021).

51. Yoon, Y., Soles, L. V. & Shi, Y. PAS-seq 2: A fast and sensitive method for global profiling of polyadenylated RNAs. Methods Enzymol. 655, 25–35 (2021).

52. Dobin, A. et al. STAR: ultrafast universal RNA-seq aligner. Bioinformatics 29, 15–21 (2013).

53. Ramírez, F., Dündar, F., Diehl, S., Grüning, B. A. & Manke, T. deepTools: a flexible platform for exploring deep-sequencing data. Nucleic Acids Res. 42, W187–191 (2014).

54. Quinlan, A. R. & Hall, I. M. BEDTools: a flexible suite of utilities for comparing genomic features. Bioinformatics 26, 841–842 (2010).

55. Wang, R., Nambiar, R., Zheng, D. & Tian, B. PolyA_DB 3 catalogs cleavage and polyadenylation sites identified by deep sequencing in multiple genomes. Nucleic Acids Res. 46, D315–D319 (2018).

56. Robinson, M. D., McCarthy, D. J. & Smyth, G. K. edgeR: a Bioconductor package for differential expression analysis of digital gene expression data. Bioinformatics 26, 139–140 (2010).

57. Tareen, A. & Kinney, J. B. Logomaker: beautiful sequence logos in Python. Bioinformatics 36, 2272–2274 (2020).

58. Linder, J., Kundaje, A. & Seelig, G. Deciphering the Impact of Genetic Variation on Human Polyadenylation. 2022.05.09.491198 Preprint at https://doi.org/10.1101/2022.05.09.491198 (2022).

59. NIH Image to ImageJ: 25 years of image analysis | Nature Methods. https://www.nature.com/articles/nmeth.2089.

60. Robinson, J. T. et al. Integrative genomics viewer. Nat. Biotechnol. 29, 24–26 (2011).

