## Supplemental Figures for "The anti-cancer compound JTE-607 reveals hidden sequence specificity of the mRNA 3′ processing machinery"

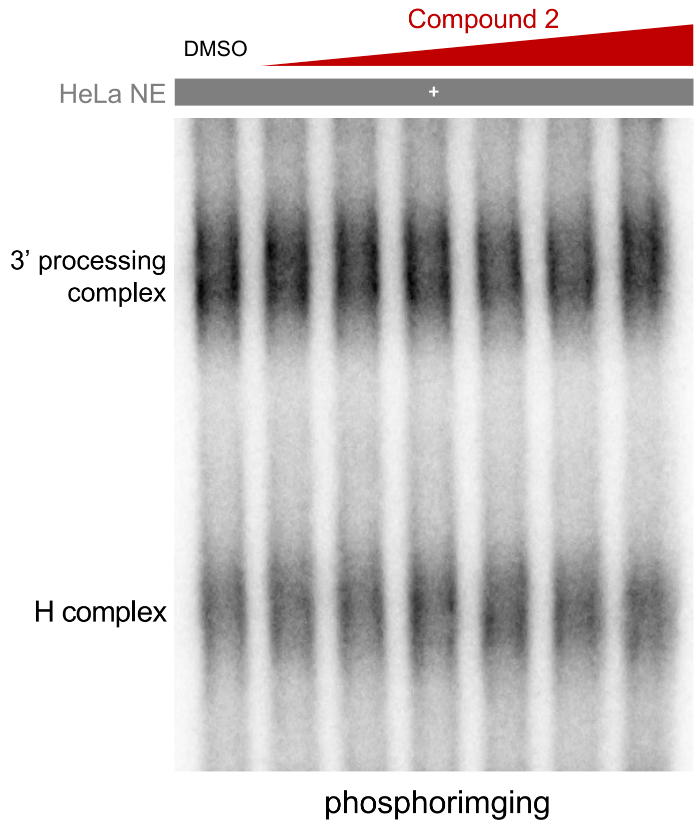


**Fig. S1. Cmp2 does not affect 3' processing complex assembly on resistant RNA.** Electrophoretic mobility shift assay (EMSA) with SVL PAS in the presence of increasing concentration of Compound 2. Same concentrations as Fig 1A and 1B were used.

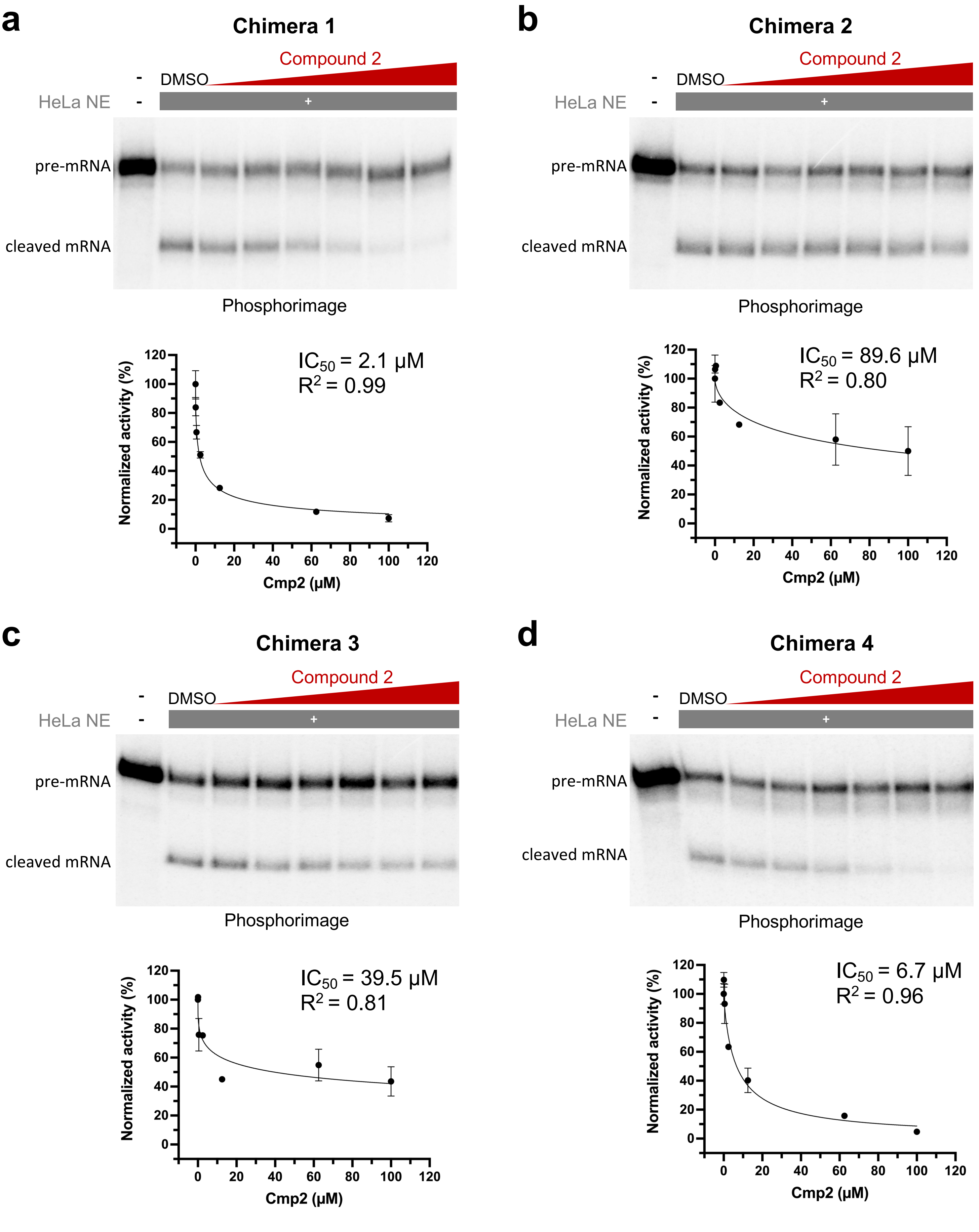


**Fig. S2. In vitro cleavage for L3 and SVL chimeras.** In vitro cleavage of L3-SVL chimeras 1-4 with increasing concentration of Compound 2 and their $IC_{50}$, similar to Fig. 1 and 2.

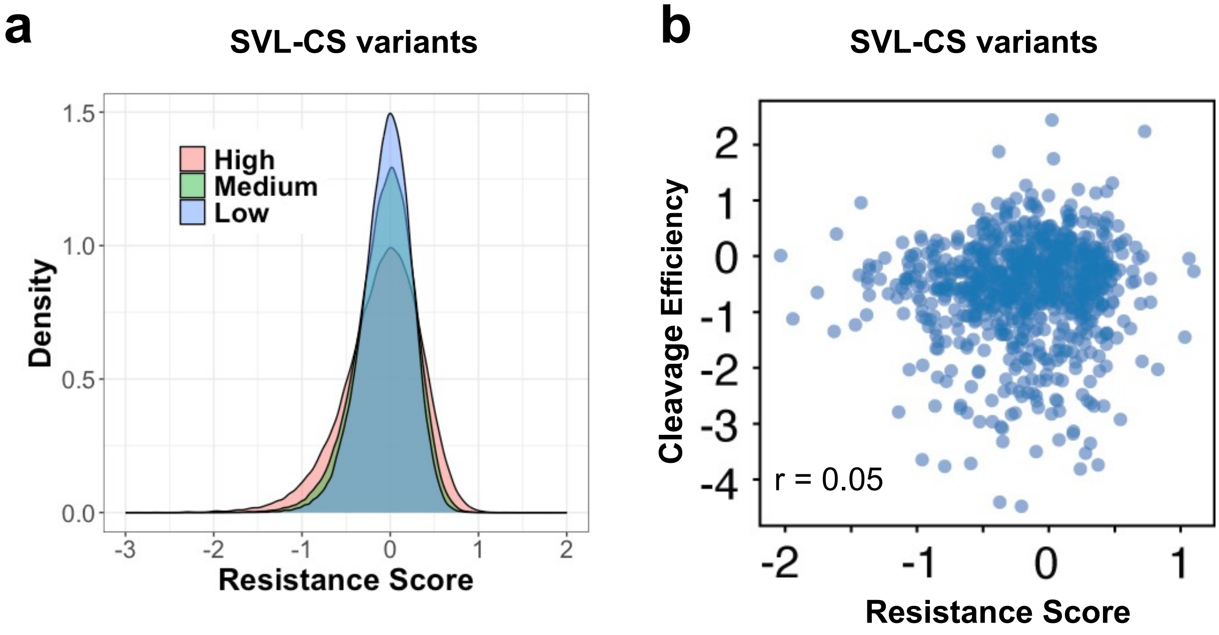


**Fig. S3. Characterization of SVL variants by MPIVA. (A)** A density plot for the resistance scores of all variants in SVL-N23 library. The low, medium, and high groups represent the screens in the presence of 0.5, 2.5, and 12.5 μM Compound 2. **(B)** A scatter plot comparing the cleavage efficiency as log(frequency in Library 2/frequency in Library 1) and the resistance score (log(frequency in Library 5/frequency in Library 2) of SVL-CS variants. Pearson correlation is shown.

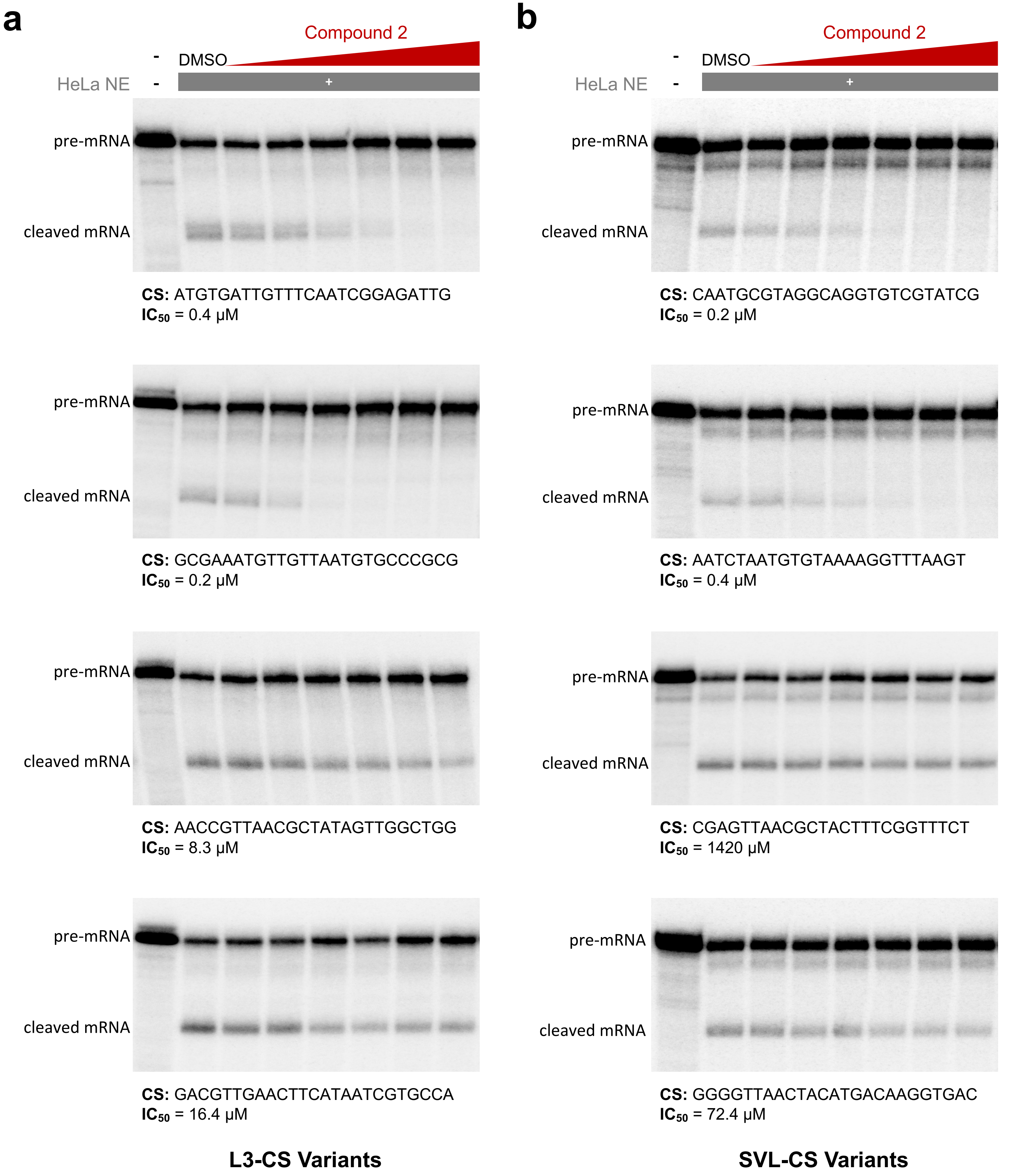


**Fig. S4. Additional validated CS variants from both backbones from in vitro MPIVA.** In vitro cleavage validation experiment of 4 more RNAs (2 sensitive and 2 resistant) from both L3- and SVL-N23 libraries. Compound 2 concentrations used are the same as Fig 1-3. The CS region sequences and their $IC_{50}$ are shown.

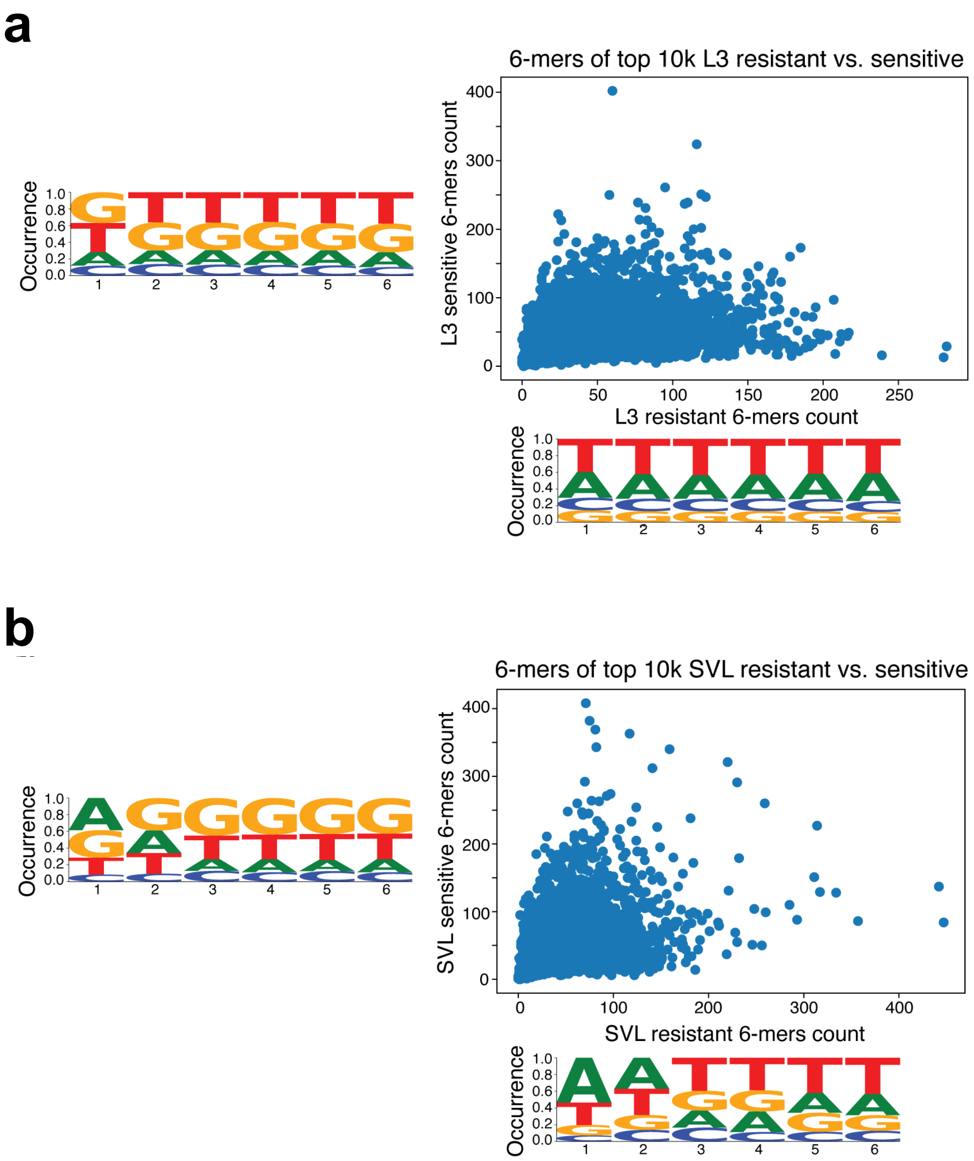


**Fig. S5. 6-mer motif analyses of top 10,000 resistant and sensitive CS variants from both backbones from the in vitro MPRA.** Counts of 6-mers in top 10k resistant and sensitive CS in **(A)** L3- and **(B)** SVL-N23 libraries backbones are plotted. The logos for 6-mers enriched in sensitive (left logo) and resistant (bottom logo) 10,000 CS variants are shown.

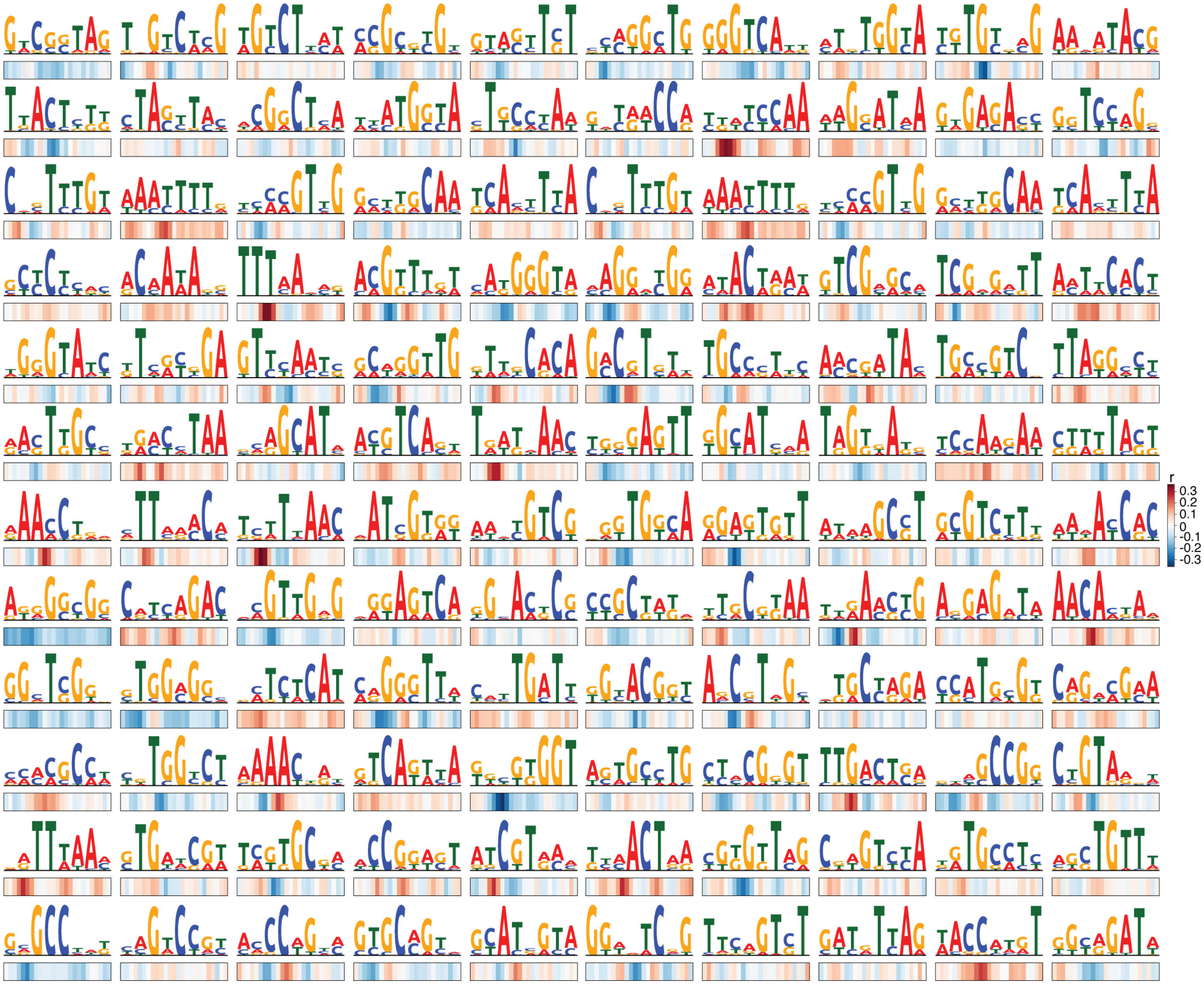


**Fig S6. C3PO layer 1 learned sequence features.** All of C3PO's layer 1 filters' max activation sequence consensus and correlations with 12.5 μM Compound 2 sensitivity predictions. Related to Fig 4D.

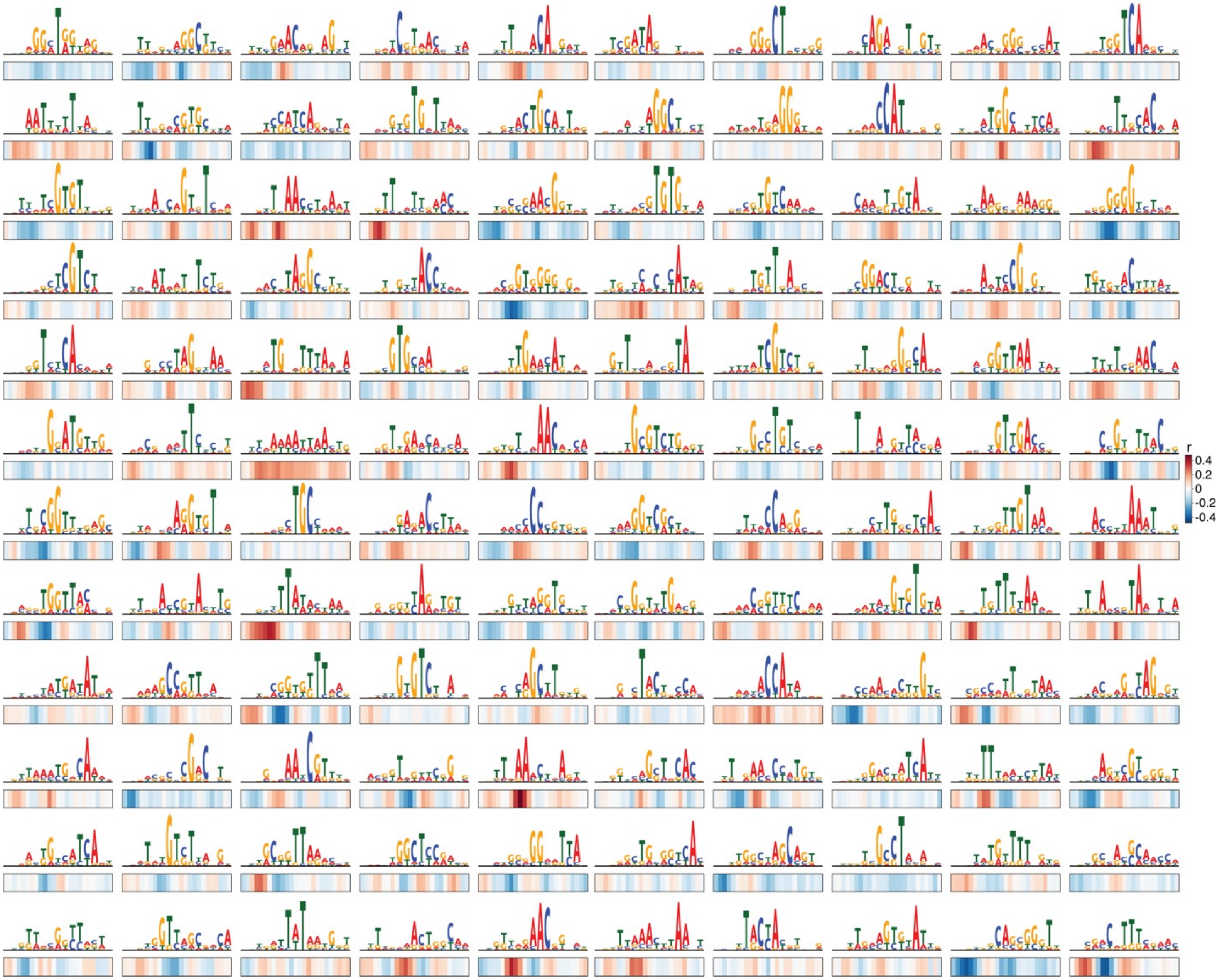


**Fig S7. C3PO layer 2 learned sequence features.** All of C3PO's layer 2 filters' max activation sequence consensus and correlations with 12.5 μM Compound 2 sensitivity predictions. Related to Figure 4E.

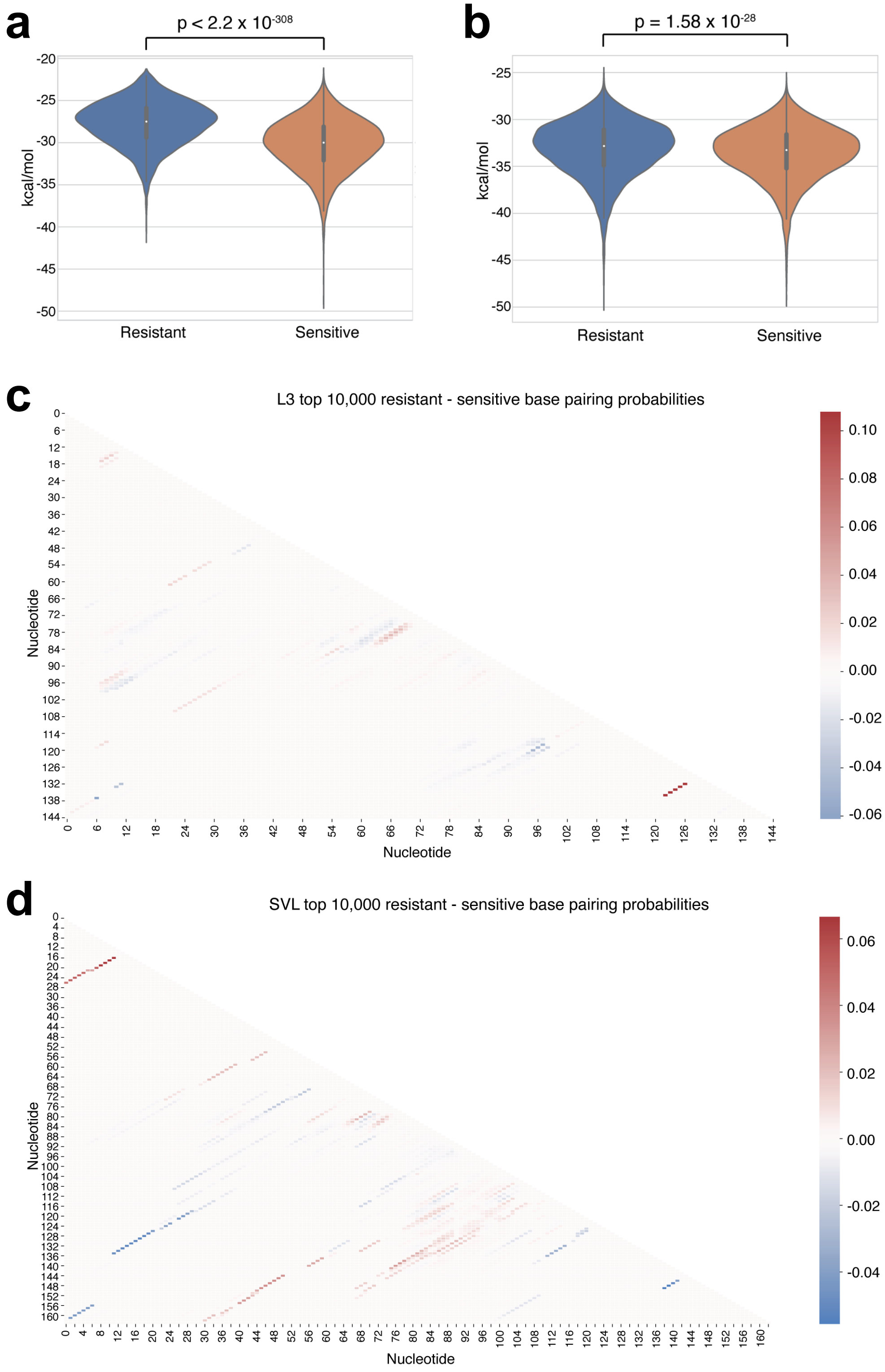


**Figure S8. ΔG of minimum free energy structures and base pairing probabilities of the top 10,000 resistant and sensitive sequences.** Comparison of minimum free energy (MFE) structures' of ΔG's from the top 10,000 resistant and sensitive **(A)** L3 and **(B)** SVL sequences. The ΔG's are significant with a p-value of < $2.2 \times 10^{-308}$ for L3 and $1.58 \times 10^{-28}$ for SVL (t-test with unequal variance). **(C)** Heatmap of the difference between top 10,000 resistant and sensitive L3 sequences' average base pairing probabilities. **(D)** Same as in panel C but for SVL.

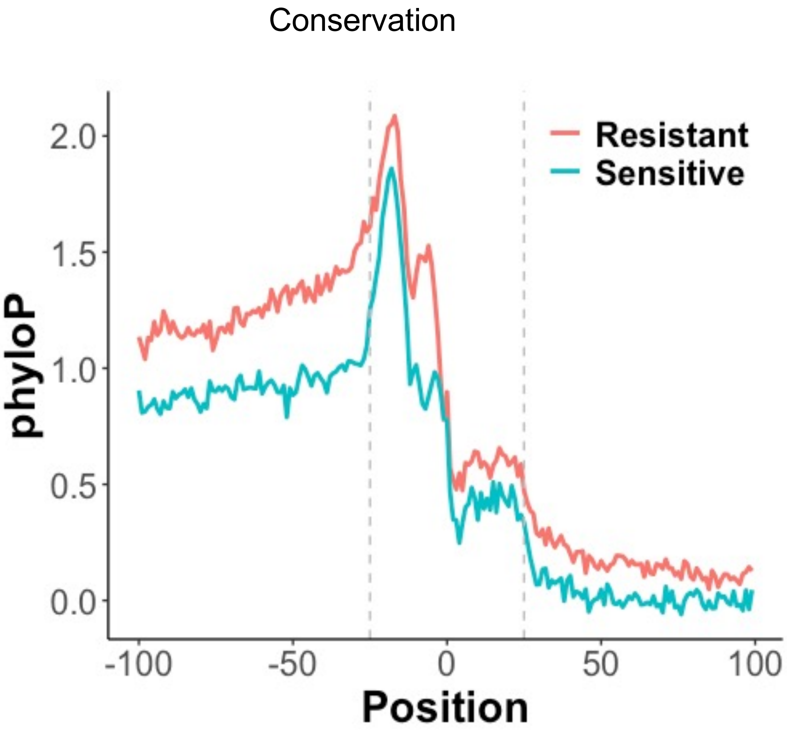


**Fig S9. Conservation of JTE-607-sensitive and –resistant PASs.** The phyloP sequence conservation score for both resistant and sensitive PASs across different species was calculated and plotted against nucleotide position of the CS. Position 0 is the YA (Y is U or C) cleavage position.
