## Supplemental Tables for "The anti-cancer compound JTE-607 reveals hidden sequence specificity of the mRNA 3′ processing machinery"

**Supplemental Table S1.** Activity and IC_50_ of all PASs plotted in Fig. 1D

| **PAS Name** | **Activity** | **IC_50_ (μM)** |
| --- | --- | --- |
| L3 | 28.04 | 0.8 |
| SVL | 36.12 | 100.2 |
| L3-SVL-Up | 51.99 | 2.2 |
| L3-SVL-Down | 54.62 | 6.7 |
| L3-SVL-CS | 44.14 | 47.8 |
| SVL-L3-Up | 31 | 39.5 |
| SVL-L3-Down | 32.46 | 89.6 |
| SVL-L3-CS | 14.2 | 0.8 |
| SVL-L3-CS2 | 13.36 | 0.6 |
| SVL-L3-CS3 | 17.67 | 0.5 |
| SVL-L3-CS4 | 9.15 | 5.5 |
| L3m3 | 4.76 | 0.7 |
| SVL230 | 10.75 | 45.2 |
| L3-SVL230-CS | 20.94 | 0.9 |
| SVL230-L3-CS | 0.15 | 20.2 |
| PerPAS | 15.04 | 1.0 |
| PerPAS-dA | 17.57 | 0.4 |
| PerPAS-GUGUm | 0.53 | 2.3 |
| PerPAS-UGUAm | 3.37 | 0.7 |
| BASP1 | 6.08 | 0.5 |
| SAU5 | 10.59 | 0.9 |
| HO-1 | 1.38 | 0.8 |
| ACTB | 19.63 | 13.9 |
| UCK2 | 5.7 | 0.5 |
| CBX6 | 2.65 | 0.3 |
| GAPDH | 38.09 | 0.5 |
| ICP27 | 8.03 | 0.5 |
| bGH | 26.08 | 38.1 |
| L3-Sen1 | 15.5 | 0.4 |
| L3-Sen52 | 5.49 | 0.2 |
| L3-Sen84 | 17.65 | 0.2 |
| L3-Rst14 | 46.65 | 42.9 |
| L3-Rst34 | 33.81 | 8.3 |
| L3-Rst52 | 41.99 | 16.4 |
| SVL-Sen3 | 9.03 | 0.2 |
| SVL-Sen38 | 18.72 | 0.6 |
| SVL-Sen127 | 10.8 | 0.4 |
| SVL-Rst303 | 18.46 | 72.4 |

**Supplemental Table S2.** R^2^ for various machine learning architectures tested

| **Model** | **Test R^2^: 0.5 µM** | **Test R^2^: 2.5 µM** | **Test R^2^: 12.5 µM** |
| --- | --- | --- | --- |
| C3PO | 0.31 | 0.552 | 0.7 |
| CNN - 4 epochs | 0.282 | 0.53 | 0.682 |
| CNN - 4 epochs, 12.5 µM only | N/A | N/A | 0.683 |
| CNN - 5 epochs | 0.301 | 0.542 | 0.687 |
| CNN - 5 epochs, 12.5 µM only | N/A | N/A | 0.689 |
| CNN - 6 epochs | 0.291 | 0.523 | 0.687 |
| CNN - 6 epochs, 12.5 µM only | N/A | N/A | 0.704 |
| CNN - 7 epochs | 0.3 | 0.543 | 0.691 |
| CNN - 7 epochs, 12.5 µM only | N/A | N/A | 0.687 |
| CNN - 8 epochs | 0.306 | 0.536 | 0.684 |
| CNN - 8 epochs, 12.5 µM only | N/A | N/A | 0.688 |
| CNN - validation early stop | 0.314 | 0.546 | 0.694 |
| CNN - validation early stop, 12.5 µM only | N/A | N/A | 0.691 |
| CNN - 4 epochs, Trial 2 | 0.311 | 0.54 | 0.688 |
| CNN - 4 epochs, 12.5 µM only, Trial 2 | N/A | N/A | 0.682 |
| CNN - 5 epochs, Trial 2 | 0.305 | 0.539 | 0.687 |
| CNN - 5 epochs, 12.5 µM only, Trial 2 | N/A | N/A | 0.695 |
| CNN - 6 epochs, Trial 2 | 0.307 | 0.548 | 0.699 |
| CNN - 6 epochs, 12.5 µM only, Trial 2 | N/A | N/A | 0.701 |
| CNN - 7 epochs, Trial 2 | 0.302 | 0.54 | 0.686 |
| CNN - 7 epochs, 12.5 µM only, Trial 2 | N/A | N/A | 0.69 |
| CNN - 8 epochs, Trial 2 | 0.296 | 0.538 | 0.69 |
| CNN - 8 epochs, 12.5 µM only, Trial 2 | N/A | N/A | 0.69 |
| CNN - validation early stop, Trial 2 | 0.307 | 0.539 | 0.688 |
| CNN - validation early stop, 12.5 µM only, Trial 2 | N/A | N/A | 0.696 |
| CNN - 6 epochs, Trial 1 out of 10 | 0.292 | 0.544 | 0.687 |
| CNN - 6 epochs, Trial 1 out of 10, 12.5 µM only | N/A | N/A | 0.701 |
| CNN - 6 epochs, Trial 2 out of 10 | 0.299 | 0.538 | 0.696 |
| CNN - 6 epochs, Trial 2 out of 10, 12.5 µM only | N/A | N/A | 0.69 |
| CNN - 6 epochs, Trial 3 out of 10 | 0.307 | 0.553 | 0.699 |
| CNN - 6 epochs, Trial 3 out of 10, 12.5 µM only | N/A | N/A | 0.701 |
| CNN - 6 epochs, Trial 4 out of 10 | 0.317 | 0.541 | 0.689 |
| CNN - 6 epochs, Trial 4 out of 10, 12.5 µM only | N/A | N/A | 0.694 |
| CNN - 6 epochs, Trial 5 out of 10 | 0.31 | 0.548 | 0.695 |
| CNN - 6 epochs, Trial 5 out of 10, 12.5 µM only | N/A | N/A | 0.69 |
| CNN - 6 epochs, Trial 6 out of 10, 12.5 µM only | N/A | N/A | 0.69 |
| CNN - 6 epochs, Trial 7 out of 10 | 0.277 | 0.514 | 0.672 |
| CNN - 6 epochs, Trial 7 out of 10, 12.5 µM only | N/A | N/A | 0.695 |
| CNN - 6 epochs, Trial 8 out of 10 | 0.304 | 0.546 | 0.693 |
| CNN - 6 epochs, Trial 8 out of 10, 12.5 µM only | N/A | N/A | 0.692 |
| CNN - 6 epochs, Trial 9 out of 10 | 0.295 | 0.545 | 0.697 |
| CNN - 6 epochs, Trial 9 out of 10, 12.5 µM only | N/A | N/A | 0.695 |
| CNN - 6 epochs, Trial 10 out of 10 | 0.305 | 0.55 | 0.697 |
| CNN - 6 epochs, Trial 10 out of 10, 12.5 µM only | N/A | N/A | 0.687 |
| CNN - hyperband training | 0.198 | 0.441 | 0.585 |
| CNN - hyperband training, 12.5 µM only | N/A | N/A | 0.586 |
| ResNet | 0.232 | 0.541 | 0.681 |
| ResNet - 7 epochs | 0.216 | 0.497 | 0.632 |
| ResNet - 12221 dilations | 0.221 | 0.494 | 0.647 |
| ResNet - 11111 dilations | 0.221 | 0.525 | 0.657 |
| ResNet - cleavage length 27 | 0.229 | 0.55 | 0.686 |
| ResNet - cleavage length 27, 75% loss for Compound 2 sensitivity predictions | 0.256 | 0.543 | 0.68 |

**Supplemental Table S3.** Log (12.5 µM/DMSO) and C3PO predicted score for all PASs plotted in Fig 4C

| **PAS Name** | **Log (12.5 µM/DMSO)** | **C3PO Predicted Score** |
| --- | --- | --- |
| L3 | -0.67 | -0.72 |
| SVL | -0.14 | 0.34 |
| BASP1 | -0.71 | 0.15 |
| SAU5 | -0.54 | 0.21 |
| PerPAS | -0.99 | -0.61 |
| PerPAS-dA | -2.13 | -0.88 |
| PerPAS-GUGUm | -0.83 | -0.60 |
| HO-1 | -0.73 | 0.15 |
| L3m3 | -1.00 | -0.81 |
| ACTB | -0.28 | 0.31 |
| UCK2 | -0.60 | 0.18 |
| CBX6 | -0.63 | -0.10 |
| GAPDH | -0.89 | 0.31 |
| ICP27 | -0.44 | -0.42 |
| bGH | -0.22 | 0.02 |
| SVL-L3-CS2 | -0.52 | 0.04 |
| SVL-L3-CS3 | -0.29 | -0.11 |
| SVL-L3-CS4 | -0.18 | 0.74 |
| L3-Sen1 | -1.62 | -1.02 |
| L3-Sen52 | -1.47 | -1.49 |
| L3-Sen84 | -1.92 | -1.92 |
| L3-Rst14 | -0.23 | 0.94 |
| L3-Rst34 | -0.26 | 0.96 |
| L3-Rst52 | -0.29 | 0.96 |
| SVL-Sen3 | -1.18 | -1.15 |
| SVL-Sen38 | -1.24 | -0.94 |
| SVL-Sen127 | -1.16 | -0.50 |
| SVL-Rst27 | -0.18 | 1.04 |
| SVL-Rst55 | -0.17 | 0.30 |
| SVL-Rst303 | -0.21 | 0.55 |

**Supplemental Table S4.** Sequence of all in vitro tested PAS in this study

| **PAS NAME** | **DNA SEQUENCE** |
| --- | --- |
| L3 | TTCTTTTTGTCACTTGAAAAACATGTAAAAATAATGTACTAGGAGACACTTTCAATAAAGGCAAATGTTTTTATTTGTACACTCTCGGGTGATTATTTACCCCCCACCCTTGCCGTCTGCGAGGTACCGAGCTC |
| SVL | ATGCTTTATTTGTGAAATTTGTGATGCTATTGCTTTATTTGTAACCATTATAAGCTGCAATAAACAAGTTAACAACAACAATTGCATTCATTTTATGTTTCAGGTTCAGGGGGAGGTGTGGGAGGTTTTTTAAAGCAAGTA |
| L3-SVL-Up | ATGCTTTATTTGTGAAATTTGTGATGCTATTGCTTTATTTGTAACCATTATAAGCTGCAATAAAGGCAAATGTTTTTATTTGTACACTCTCGGGTGATTATTTACCCCCCACCCTTGCCGTCTGCGA |
| L3-SVL-Down | TTCTTTTTGTCACTTGAAAAACATGTAAAAATAATGTACTAGGAGACACTTTCAATAAAGGCAAATGTTTTTATTTGTACATTCATTTTATGTTTCAGGTTCAGGGGGAGGTGTGGGAGGTTTTTTAAAGCAAGTA |
| L3-SVL-CS | TTCTTTTTGTCACTTGAAAAACATGTAAAAATAATGTACTAGGAGACACTTTCAATAAACAAGTTAACAACAACAATTGCACTCTCGGGTGATTATTTACCCCCCACCCTTGCCGTCTGCGA |
| SVL-L3-Up | TTCTTTTTGTCACTTGAAAAACATGTAAAAATAATGTACTAGGAGACACTTTCAATAAACAAGTTAACAACAACAATTGCATTCATTTTATGTTTCAGGTTCAGGGGGAGGTGTGGGAGGTTTTTTAAAGCAAGTA |
| SVL-L3-Down | ATGCTTTATTTGTGAAATTTGTGATGCTATTGCTTTATTTGTAACCATTATAAGCTGCAATAAACAAGTTAACAACAACAATTGCACTCTCGGGTGATTATTTACCCCCCACCCTTGCCGTCTGCGA |
| SVL-L3-CS | ATGCTTTATTTGTGAAATTTGTGATGCTATTGCTTTATTTGTAACCATTATAAGCTGCAATAAAGGCAAATGTTTTTATTTGTACATTCATTTTATGTTTCAGGTTCAGGGGGAGGTGTGGGAGGTTTTTTAAAGCAAGTA |
| SVL-L3-CS2 | ATGCTTTATTTGTGAAATTTGTGATGCTATTGCTTTATTTGTAACCATTATAAGCTGCAATAAACAAGTTAACAACAATTTGTACATTCATTTTATGTTTCAGGTTCAGGGGGAGGTGTGGGAGGTTTTTTAAAGCAAGTA |
| SVL-L3-CS3 | ATGCTTTATTTGTGAAATTTGTGATGCTATTGCTTTATTTGTAACCATTATAAGCTGCAATAAACAAGTTTGTTTTTAACAATTGCATTCATTTTATGTTTCAGGTTCAGGGGGAGGTGTGGGAGGTTTTTTAAAGCAAGTA |
| SVL-L3-CS4 | ATGCTTTATTTGTGAAATTTGTGATGCTATTGCTTTATTTGTAACCATTATAAGCTGCAATAAACAAGTTAACATTTATTTGCATTCATTTTATGTTTCAGGTTCAGGGGGAGGTGTGGGAGGTTTTTTAAAGCAAGTA |
| L3m3 | TTCTTTTTGTCACTTGAAAAACATGGGAAAATAATGGGCTAGGAGACACTTTCAATAAAGGCAAATGTTTTTATTTTTACACTCTCGGGTGATTATTTACCCCCCACCCTTGCCGTCTGCGAGGTACCGAGCTC |
| SVL230 | AGACATGATAAGATACATTGATGAGTTTGGACAAACCACAACTAGAATGCAGTGAAAAAAATGCTTTATTTGTGAAATTTGTGATGCTATTGCTTTATTTGTAACCATTATAAGCTGCAATAAACAAGTTAACAACAACAATTGCATTCATTTTATGTTTCAGGTTCAGGGGGAGGTGTGGGAGGTTTTTTAAAGCAAGTAAAACCTCCAGATCCCCGGGTACCGAGCTC |
| L3-SVL230-CS | TTCTTTTTGTCACTTGAAAAACATGTAAAAATAATGTACTAGGAGACACTTTCAATAAAGGCAAAAACAACAACAATTGCACTCTCGGGTGATTATTTACCCCCCACCCTTGCCGTCTGCGAGGTACCGAGCTC |
| SVL230-L3-CS | AGACATGATAAGATACATTGATGAGTTTGGACAAACCACAACTAGAATGCAGTGAAAAAAATGCTTTATTTGTGAAATTTGTGATGCTATTGCTTTATTTGTAACCATTATAAGCTGCAATAAACAAGTTTGTTTTTATTTGTACATTCATTTTATGTTTCAGGTTCAGGGGGAGGTGTGGGAGGTTTTTTAAAGCAAGTAAAACCTCCAGATCCCCGGGTACCGAGCTC |
| PerPAS | TTTTTTTTTTTGTAAATTAATTTTTAATAAAGTTGTTTTTTACACGTTGTCTTGTGTGTCGTCTTTTTTTTCAGC |
| PerPAS-dA | TTTTTTTTTTTGTAAATTAATTTTTAATAAAGTTGTTTTTTGCGCGTTGTCTTGTGTGTCGTCTTTTTTTTCAGC |
| PerPAS-GUGUm | TTTTTTTTTTTGTAAATTAATTTTTAATAAAGTTGTTTTTTACACGTTGTCTTCTCTCTCGTCTTTTTTTTCAGC |
| PerPAS-UGUAm | TTTTTTTTTTTCTACATTAATTTTTAATAAAGTTGTTTTTTACACGTTGTCTTGTGTGTCGTCTTTTTTTTCAGC |
| BASP1 | TGAAAGGGAAAGCCTAGCAAGTCTATTAATAAGCTCACTTCCCATTTATCCCAGTGTACCTGGAGCATTAAGCTAAGACGTTCATCCACAGGCTTAAAAACTTACATCAAGCACTACTGAACTTTACAAGCTGGAATAAACAATGCCTACTAAATAAAAGATTTATAAAATTGTTCTGTCTTATTTTTGTGATCTCTTGTAAATGTTTTTTTTTTTTTTTTTTTTAAATATCCAAAGAAGACCTGTGAACTATTATTTGTCAGAAGCAATTGCCCTTGGTATCTGATTCTGTTGAAAGAA |
| SAU5 | GCCGCCCAACCCGAGCGACCTTCCCCTCCCACTTCCCCCCCCCTACACACCAACTCCGCCCTCGCCGTCTTGGCCGTGCGCGGCCCCGTGCGTCCGTCTCAATAAAGCCAGGTTAAATCCGTGACGTGGTGTGTTTGGCGTGTGTCTCTGAAATGGCGGAAACCGACATGCAAATGGGATTCATGGACATGTTACACCCCCCTGAC |
| HO-1 | GGCACTGTGGCCTTGGTCTAACTTTTGTGTGAAATAATAAACAACATTGTCTGATAGTAGCTTGAAGTAGTTTTCATGGGCTTTGTTATTCTTGGGGAACTGACCTTTTCCTCCCTGGTTTCTTGCGTGCTCGGTAGGA |
| ACTB | TTTTTTGTCCCCCAACTTGAGATGTATGAAGGCTTTTGGTCTCCCTGGGAGTGGGTGGAGGCAGCCAGGGCTTACCTGTACACTGACTTGAGACCAGTTGAATAAAAGTGCACACCTTAAAAATGAGGCCAAGTGTGACTTTGTGGTGTGGCTGGGTTGGGGGCAGCAGAGGGTGAACCCTGCAGGAGGGTGAACCCTGCAAAAGGGTGGGGCAGTGGGGGCCAAC |
| UCK2 | CCCCCCTTTTAAGATGCTTGCTCCTCTCCCTTTTCTTTTTACCACCCTACCTTTATTGTTAGTGGTTACAAAGTGACCACATATTATGTACTTTGCTGTAAATAAAGACAGACAAAAAGGCTCTCGCCTTCTGTGTGATGCTTGGCCCAGAGCAGCGACCGAATCCTGGCTGTGTGGCCCAAGTGGCTCAGGAAGGGCCATGCTGTGCATGTGTGGTGTAGA |
| CBX6 | GGGTCTGTGCCGATTACTCTGTCTTGTACGTTTGTTCTGCTGCTCTTCAATATTGTATCAACGCCAGGAAAGGGGGGTGAAAAGCCTCTTTTACCCCCCAAATAAATTGTCACATTCCGAAGCTGAGGCCTAGCCCCTAGGTTGGGGTGTGTCTGTGTCTTCTTCCAGCTGTGACTGGCTTTTCAAAAGTAGCAGGCCCATGTCCCTCCAGTGACAGGTGAAGAGGGG |
| GAPDH | TCAGTCCCCCACCACACTGAATCTCCCCTCCTCACAGTTGCCATGTAGACCCCTTGAAGAGGGGAGGGGCCTAGGGAGCCGCACCTTGTCATGTACCATCAATAAAGTACCCTGTGCTCAACCAGTTACTTGTCCTGTCTTATTCTAGGGTCTGGGGCAGAGGGGAGGGAAGCTGGGCTTGTGTCAAGGTGAGACATTCTTGCTGGGGAGGGACCTGGTATGT |
| ICP27 | GTGTTCGAGTCGTGTCTGCGAGTTGACGGCCAGTCACATCGTCGCCCCCCCGTACGTGCACGGCAAATATTTTTATTGCAACTCCCTGTTTTAGGTACAATAAAAACAAAACATTTCAAACAAATCGCCCCTCGTGTTGTCCTTCTTTGCTCATGGCCGGCGGGGCGTGGGTCACGGCAGATGGCGGGGGTGGGCCCGGCG |
| bGH | CTGTGCCTTCTAGTTGCCAGCCATCTGTTGTTTGCCCCTCCCCCGTGCCTTCCTTGACCCTGGAAGGTGCCACTCCCACTGTCCTTTCCTAATAAAATGAGGAAATTGCATCGCATTGTCTGAGTAGGTGTCATTCTATTCTGGGGGGTGGGGTGGGGCAGGACAGCAAGGGGGAGGATTGGGAAGACAATAGCAGGCATGCTGGGGATGCGGTGGGCTCTATGG |
| L3-Sen1 | TTCTTTTTGTCACTTGAAAAACATGTAAAAATAATGTACTAGGAGACACTTTCAATAAAATGTGATTGTTTCAATCGGAGATTGTCGGGTGATTATTTACCCCCCACCCTTGCCGTCTGCGAGGTACCGAGCTC |
| L3-Sen52 | TTCTTTTTGTCACTTGAAAAACATGTAAAAATAATGTACTAGGAGACACTTTCAATAAACAATGTGCTGTTCAAAGGCGGTGGCTCGGGTGATTATTTACCCCCCACCCTTGCCGTCTGCGAGGTACCGAGCTC |
| L3-Sen84 | TTCTTTTTGTCACTTGAAAAACATGTAAAAATAATGTACTAGGAGACACTTTCAATAAAGCGAAATGTTGTTAATGTGCCCGCGTCGGGTGATTATTTACCCCCCACCCTTGCCGTCTGCGAGGTACCGAGCTC |
| L3-Rst14 | TTCTTTTTGTCACTTGAAAAACATGTAAAAATAATGTACTAGGAGACACTTTCAATAAAAAGGTTAACGCTCATATGGTTCGTTTCGGGTGATTATTTACCCCCCACCCTTGCCGTCTGCGAGGTACCGAGCTC |
| L3-Rst34 | TTCTTTTTGTCACTTGAAAAACATGTAAAAATAATGTACTAGGAGACACTTTCAATAAAAACCGTTAACGCTATAGTTGGCTGGTCGGGTGATTATTTACCCCCCACCCTTGCCGTCTGCGAGGTACCGAGCTC |
| L3-Rst52 | TTCTTTTTGTCACTTGAAAAACATGTAAAAATAATGTACTAGGAGACACTTTCAATAAAGACGTTGAACTTCATAATCGTGCCATCGGGTGATTATTTACCCCCCACCCTTGCCGTCTGCGAGGTACCGAGCTC |
| SVL-Sen3 | ATGCTTTATTTGTGAAATTTGTGATGCTATTGCTTTATTTGTAACCATTATAAGCTGCAATAAACAATGCGTAGGCAGGTGTCGTATCGATTTTATGTTTCAGGTTCAGGGGGAGGTGTGGGAGGTTTTTTAAAGCAAGTA |
| SVL-Sen38 | ATGCTTTATTTGTGAAATTTGTGATGCTATTGCTTTATTTGTAACCATTATAAGCTGCAATAAAAACATGTCGTGCATTTGTTTCATTGATTTTATGTTTCAGGTTCAGGGGGAGGTGTGGGAGGTTTTTTAAAGCAAGTA |
| SVL-Sen127 | ATGCTTTATTTGTGAAATTTGTGATGCTATTGCTTTATTTGTAACCATTATAAGCTGCAATAAAAATCTAATGTGTAAAAGGTTTAAGTATTTTATGTTTCAGGTTCAGGGGGAGGTGTGGGAGGTTTTTTAAAGCAAGTA |
| SVL-Rst27 | ATGCTTTATTTGTGAAATTTGTGATGCTATTGCTTTATTTGTAACCATTATAAGCTGCAATAAACGAGTTAACGCTACTTTCGGTTTCTATTTTATGTTTCAGGTTCAGGGGGAGGTGTGGGAGGTTTTTTAAAGCAAGTA |
| SVL-Rst55 | ATGCTTTATTTGTGAAATTTGTGATGCTATTGCTTTATTTGTAACCATTATAAGCTGCAATAAAGTGCGGTAACGCAGAATTTTGTAATATTTTATGTTTCAGGTTCAGGGGGAGGTGTGGGAGGTTTTTTAAAGCAAGTA |
| SVL-Rst303 | ATGCTTTATTTGTGAAATTTGTGATGCTATTGCTTTATTTGTAACCATTATAAGCTGCAATAAAGGGGTTAACTACATGACAAGGTGACATTTTATGTTTCAGGTTCAGGGGGAGGTGTGGGAGGTTTTTTAAAGCAAGTA |

**Supplemental Table S5.** Oligonucleotides and synthesized DNA used in this study

| **OLIGO NAME** | **DNA SEQUENCE** |
| --- | --- |
| **For PAS cloning** | |
| L3-F | ACAGGATCCTTCTTTTTGTCACTTGAAAAACATGTA |
| L3-R | ACAGAATTCTCGCAGACGGCAAG |
| SVL230-F | ACAGGATCCAGACATGATAAGATACATTGATGAG |
| SVL230-R | ACAGAATTCGAGCTCGGTACCCG |
| SVL-F | ACAGGATCCATGCTTTATTTGTGAAATTTGTGATGCTATTG |
| SVL-R | ACAGAATTCTACTTGCTTTAAAAAACCTCCCACAC |
| L3-SVL Up-F | GTAACCATTATAAGCTGCAATAAAGGCAAATGTTTTTA |
| L3-SVL Up-R | TAAAAACATTTGCCTTTATTGCAGCTTATAATGGTTAC |
| L3-SVL Down-F | CAAATGTTTTTATTTGTACATTCATTTTATGTTTCA |
| L3-SVL Down-R | TGAAACATAAAATGAATGTACAAATAAAAACATTTG |
| SVL-L3 Up-F | GTACTAGGAGACACTTTCAATAAACAAGTTAACAAC |
| SVL-L Up-R | GTTGTTAACTTGTTTATTGAAAGTGTCTCCTAGTAC |
| SVL-L3 Down-F | GTTAACAACAACAATTGCACTCTCGGGTGATTATTTA |
| SVL-L3 Down-R | TAAATAATCACCCGAGAGTGCAATTGTTGTTGTTAAC |
| L3-SVL CS-F | CAAGTTAACAACAACAATTGCACTCTCGGGTGATTATTTAC |
| L3-SVL CS-R | CAATTGTTGTTGTTAACTTGTTTATTGAAAGTGTCTCC |
| SVL-L3 CS-F | GGCAAATGTTTTTATTTGTACATTCATTTTATGTTTCAG |
| SVL-L3 CS-R | TACAAATAAAAACATTTGCCTTTATTGCAGCTTATAATG |
| SVL-L3 CS2-F | ATTTGTACATTCATTTTATGTTTCAGGTTCAG |
| SVL-L3 CS2-R | TGTTGTTAACTTGTTTATTGCAGC |
| SVL-L3 CS3-F | TGTTTTTAACAATTGCATTCATTTTATGTTTCAG |
| SVL-L3 CS3-R | AACTTGTTTATTGCAGCTTATAATG |
| SVL-L3 CS4-F | TTTATTTGCATTCATTTTATGTTTCAGGTTCAG |
| SVL-L3 CS4-R | TGTTAACTTGTTTATTGCAGC |
| L3m3-F | CAAATGTTTTTATTTTTACACTCTCGGGTG |
| L3m3-R | CACCCGAGAGTGTAAAAATAAAAACATTTG |
| PerPAS-F | ACATCTAGATTTTTTTTTTTGTAAATTAATTTTTAATAAAGTTGTTTTTT |
| PerPAS-R | ACACTCGAGGCTGAAAAAAAAGACGACACACAAGACAACGTGTAAAAAACAACTTT |
| PerPAS-mut-F | ACATCTAGATTTTTTTTTTTGTAAATTAATTTTTAACAAAGTTGTTTTTT |
| PerPAS-dA-R | ACACTCGAGGCTGAAAAAAAAGACGACACACAAGACAACGCGCAAAAAACAACTTT |
| PerPAS-UGUAmut-R | ACATCTAGATTTTTTTTTTTCTACATTAATTTTTAATAAAGTTGTTTTTT |
| PerPAS-GUGUmut-R | ACACTCGAGGCTGAAAAAAAAGACGAGAGAGAAGACAACGTGTAAAAAACAACTTT |
| BASP1-F | AGTCTAGATGAAAGGGAAAGCCTAGC |
| BASP1-R | TTCTCGAGTTCTTTCAACAGAATCAG |
| SAU5-F | ACATCTAGAGCCGCCCAACCCGAGCGACCT |
| SAU5-R | ACACTCGAGGTCAGGGGGGTGTAACATGTCCA |
| HO1-F | ACTCTAGAGGCACTGTGGCCTTGGTCTAA |
| HO1-R | ATCTCGAGtCCTACCGAGCACGCAAGAA |
| ACTB-F | ACATCTAGATTTTTTGTCCCCCAACTTGAG |
| ACTB-R | ACACTCGAGGTTGGCCCCCACTGCCCCAC |
| UCK2-F | ACATCTAGACCCCCCTTTTAAGATGCTTG |
| UCK2-R | ACACTCGAGTCTACACCACACATGCACAG |
| CBX6-F | ACATCTAGAGGGTCTGTGCCGATTACTCT |
| CBX6-R | ACACTCGAGCCCCTCTTCACCTGTCACTG |
| GAPDH-F | ACATCTAGATCAGTCCCCCACCACACTGA |
| GAPDH-R | ACACTCGAGACATACCAGGTCCCTCCCCA |
| ICP27-F | ACATCTAGAGTGTTCGAGTCGTGTCTGCGAG |
| ICP27-R | ACACTCGAGCGCCGGGCCCACCCCCGCCATCTGCCGTGACCCAC |
| bGH-F | ACAGGATCCCTGTGCCTTCTAGTTGCCAG |
| bGH-R | ACAGAATTCCCATAGAGCCCACCGCATC |
| L3 Sen1-F | ATGTGATTGTTTCAATCGGAGATTGTCGGGTGATTATTTACCCCCCAC |
| L3 Sen52-F | CAATGTGCTGTTCAAAGGCGGTGGCTCGGGTGATTATTTACCCCCCAC |
| L3 Sen84-F | GCGAAATGTTGTTAATGTGCCCGCGTCGGGTGATTATTTACCCCCCAC |
| L3 Rst14-F | AAGGTTAACGCTCATATGGTTCGTTTCGGGTGATTATTTACCCCCCAC |
| L3 Rst34-F | AACCGTTAACGCTATAGTTGGCTGGTCGGGTGATTATTTACCCCCCAC |
| L3 Rst52-F | GACGTTGAACTTCATAATCGTGCCATCGGGTGATTATTTACCCCCCAC |
| L3 linear-R | TTTATTGAAAGTGTCTCCTAGTACATTATTTTTAC |
| SVL Sen3-F | CAATGCGTAGGCAGGTGTCGTATCGATTTTATGTTTCAGGTTCAGGGGGAG |
| SVL Sen38-F | AACATGTCGTGCATTTGTTTCATTGATTTTATGTTTCAGGTTCAGGGGGAG |
| SVL Sen127-F | AATCTAATGTGTAAAAGGTTTAAGTATTTTATGTTTCAGGTTCAGGGGGAG |
| SVL Rst27-F | CGAGTTAACGCTACTTTCGGTTTCTATTTTATGTTTCAGGTTCAGGGGGAG |
| SVL Rst55-F | GTGCGGTAACGCAGAATTTTGTAATATTTTATGTTTCAGGTTCAGGGGGAG |
| SVL Rst303-F | GGGGTTAACTACATGACAAGGTGACATTTTATGTTTCAGGTTCAGGGGGAG |
| SVLst linear-R | TTTATTGCAGCTTATAATGGTTACAAATAAAGC |
| **For MPIVA N23 screen** | |
| T7-L3 N23 | TAATACGACTCACTATAGGGATAATTTCTTTTTGTCACTTGAAAAACATGTAAAAATAATGTACTAGGAGACACTTTCAATAAANNNNNNNNNNNNYANNNNNNNNNNNTCGGGTGATTATTTACCCCCCACCCTTGCCGTCTGCGA |
| T7-SVL N23 | TAATACGACTCACTATAGGGATAATATGCTTTATTTGTGAAATTTGTGATGCTATTGCTTTATTTGTAACCATTATAAGCTGCAATAAANNNNNNNNNNNYANNNNNNNNNNNNATTTTATGTTTCAGGTTCAGGGGGAGGTGTGGGAGGTTTTTTAAAGCAAGTA |
| L3 N23 Gibson-F | CTATAGGGCGAATTGGAGCTCTTCTTTTTGTCACTTGAAAAACATGTA |
| L3 N23 Gibson-R | GTATCGATAAGCTTGATATCGAATTCTCGCAGACGGCAAGG |
| SVL N23 Gibson-F | CTATAGGGCGAATTGGAGCTCATGCTTTATTTGTGAAATTTGTGATGC |
| SVL N23 Gibson-R | GTATCGATAAGCTTGATATCGAATTCTACTTGCTTTAAAAAACCTCCCACAC |
